# SKOOTS: Skeleton-oriented object segmentation for mitochondria

**DOI:** 10.1101/2023.05.05.539611

**Authors:** Christopher J Buswinka, Richard T. Osgood, Hidetomi Nitta, Artur A. Indzhykulian

## Abstract

Segmenting individual instances of mitochondria from imaging datasets can provide rich quantitative information, but is prohibitively time-consuming when done manually, prompting interest in the development of automated algorithms using deep neural networks. Existing solutions for various segmentation tasks are optimized for either: high-resolution three-dimensional imaging, relying on well-defined object boundaries (e.g., whole neuron segmentation in volumetric electron microscopy datasets); or low-resolution two-dimensional imaging, boundary-invariant but poorly suited to large 3D objects (e.g., whole-cell segmentation of light microscopy images). Mitochondria in whole-cell 3D electron microscopy datasets often lie in the middle ground—large, yet with ambiguous borders, challenging current segmentation tools. To address this, we developed **sk**eleton-**o**riented **o**bjec**t s**egmentation (SKOOTS)—a novel approach that efficiently segments large, densely packed mitochondria. SKOOTS accurately and efficiently segments mitochondria in previously difficult contexts and can also be applied to segment other objects in 3D light microscopy datasets. This approach bridges a critical gap between existing segmentation approaches, improving the utility of automated analysis of three-dimensional biomedical imaging data. We demonstrate the utility of SKOOTS by applying it to segment over 15,000 cochlear hair cell mitochondria across experimental conditions in under 2 hours on a consumer-grade PC, enabling downstream morphological analysis that revealed subtle structural changes following aminoglycoside exposure—differences not detectable using analysis approaches currently used in the field.

## INTRODUCTION

Mitochondrial dynamics, driven by processes such as fission and fusion that regulate their size and number, are thought to play important roles in the pathology of numerous diseases^1^. Mitochondrial dysfunction has been implicated in Alzheimer’s^2,3^, Huntington’s^4^, Parkinson’s^5^, and cardiovascular diseases^6^, as well as cancer, auditory neuropathy^7,8^, and many other conditions. Mitochondria are abundant in presynaptic terminals^9,10^, help regulate Ca^2+^ dynamics^11-13^, and participate in apoptotic signaling cascades^14-17^. In metabolically active cells, they can occupy a significant fraction of the total cellular volume^18^; however, their relatively small size and variable morphology make resolving single instances of individual mitochondria, and performing automated image analysis, challenging^19^.

Although recent advances in light microscopy have enabled the visualization of individual mitochondria in some cases^20^, three-dimensional electron microscopy (3D EM) techniques such as focused ion beam scanning electron microscopy (FIB-SEM)^21^ or serial blockface SEM offer nanometer-scale resolution and are better suited to capture subtle morphological features. These techniques, when used to image entire or multiple cells, often generate large and rich datasets over large spatial domains^22^. To assess mitochondrial volume, morphology, and spatial distribution, and how these features change under experimental conditions, accurate instance segmentation of each mitochondrion is essential. However, analyzing thousands of mitochondria at high resolution manually is impractical and presents a significant bottleneck.

Reliable automated segmentation of mitochondria in 3D EM datasets remains a persistent challenge^19^. In fields such as connectomics, where the goal is to reconstruct neural circuits from high-resolution 3D EM datasets, segmentation must be nearly flawless across vast imaging volumes^23^. This demand has spurred the development of three-dimensional algorithms, most of which predict object boundaries by generating affinity graphs^24-26^. In these approaches, a network predicts the likelihood that adjacent 3D pixels (voxels) belong to the same object^24^. Voxels with high mutual affinity are grouped into connected components via watershed segmentation, enabling inference of inter-voxel boundaries^27-31^.

When affinity predictions are accurate, these methods are fast and effective^32^. However, segmentation performance deteriorates in cases where object boundaries are ambiguous, for example when mitochondria are densely packed and in contact with one another^30^. These conditions can result in poor affinity prediction and under-segmentation, where multiple mitochondria are erroneously merged into a single segmentation mask.

Ambiguous boundaries between densely packed objects are not unique to EM; similar challenges arise in fluorescence microscopy of densely labeled cells. Approaches like *Cellpose*^33^ and its successor *Omnipose*^34^, the two most widely used tools for 2D light microscopy segmentation, address this issue by predicting optical flows—vectors that direct pixels toward the center of individual objects, effectively separating touching cells. Optical flow-based methods rely on pixels being directed toward meaningful targets within each object. In *Cellpose*, pixels are guided along a diffusion gradient toward the cell’s center of mass. However, in some cases, such as in curled bacteria^34^, the center of mass may lie outside the cell, limiting segmentation accuracy. This limitation led to the core innovation in *Omnipose*: pixels flow along the gradient of an Eikonal distance function^35^, which points toward the local center of mass regardless of shape. Applying these flows generates clusters of pixels corresponding to individual instances, which must then be identified using a clustering algorithm, typically *DBSCAN*^36^.

These approaches are well suited for two-dimensional biomedical instance segmentation. Although they can be extended to 3D, they are constrained in practice by the clustering step: *DBSCAN* scales poorly in memory and slows exponentially in time [O(N^2^)], which becomes prohibitive when applied to volumetric datasets^37^. For instance, *Omnipose* is reportedly limited to processing a 78(x) × 78(y) × 78(z) voxel cube on a 12 GB consumer-grade GPU^34^. Segmenting larger volumes therefore requires either downscaling, which compromises resolution, or computationally expensive tiling, or both, making such approaches unsuitable for segmenting mitochondria in high-resolution 3D EM datasets.

While these limitations are less pronounced in 2D, they can be partially circumvented by applying 2D segmentation models across multiple planes of a 3D image and reconstructing a volumetric mask. This workaround is employed by *Cellpose* and *Empanada*^38^, a more recent mitochondria segmentation platform. However, the reconstruction step introduces an additional point of failure and may propagate segmentation errors.

Additional challenges limit the development and implementation of 3D machine learning segmentation approaches. Training data are time-consuming to generate, and existing datasets are often inadequate for specific analysis tasks. Moreover, practical implementation is constrained by computational efficiency and hardware requirements. For broad accessibility, a 3D segmentation method must be not only accurate but also efficient. Approaches that require high-performance computing are less likely to be adopted than those that run on a standard personal computer. Similarly, methods that execute slowly are less practical than those that deliver results in near real time. In this work, we take steps to address these limitations.

In this study, we present a new approach for instance segmentation of mitochondria in high-resolution 3D EM datasets: **s**keleton-**o**riented **o**bjec**t s**egmentation (SKOOTS). We demonstrate that SKOOTS is fast, efficient, and accurate—even for densely packed mitochondria with diverse morphologies. To train and validate the algorithm, we created a FIB-SEM dataset for which we manually annotated mitochondria instance masks. We selected auditory hair cells as our model system due to their exceptional metabolic demands and dense mitochondrial content. SKOOTS was trained using a variant of the U-Net deep learning architecture^39^, modified to improve accuracy per parameter compared to conventional implementations^40^. We compare SKOOTS with several existing mitochondria segmentation approaches and demonstrate superior performance in both accuracy and computational efficiency, particularly for densely packed structures. We the utility of SKOOTS in biomedical research by applying it to a large-scale analysis of cochlear hair cell mitochondria following aminoglycoside treatment—a widely used model for studying drug-induced hearing loss. Using SKOOTS, we segmented and analyzed over 15,000 mitochondria from FIB-SEM datasets of treated and control cells, revealing statistically significant morphological changes that were not detectable using existing analysis pipelines. These findings illustrate the power of our approach to uncover subtle structural alterations that may contribute to cellular vulnerability to ototoxic damage. Across all datasets and experimental conditions SKOOTS demonstrated both scalability and robustness. Finally, we show that SKOOTS is a generalizable algorithm that can be readily applied to 3D instance segmentation of diverse biological structures in high-resolution light microscopy datasets, including densely packed cytoplasmic compartments and plant tissues—underscoring its broad utility across imaging modalities.

## RESULTS

### Skeleton embedding allows for segmentation of mitochondria

The key steps of the SKOOTS segmentation are illustrated in **Figure 1**. Densely packed mitochondria sometimes exhibit unclear boundaries, making affinity graph predictions unreliable and resulting in poor segmentation outcomes. In whole-cell FIB-SEM datasets, they are also frequently imaged at resolutions too high for existing 3D optical flow techniques to handle efficiently. Our method—**sk**eleton **o**riented **o**bjec**t s**egmentation (SKOOTS)— addresses these challenges by leveraging object skeletons. Object skeletons are defined as connected paths through local centers of mass^41^. While mitochondria may be densely packed and in direct membrane contact, their skeletons will remain distinct. This approach addresses a critical limitation of some optical flow-based methods: determining where to direct flow when the model cannot “see” the entire object. Furthermore, we explicitly predict object skeletons as a 3D volume. As a result, unique ID labels can be efficiently assigned using a flood fill algorithm^42^—an approach that is more computationally efficient than the clustering algorithms typically used for this task.

**Figure 1.**
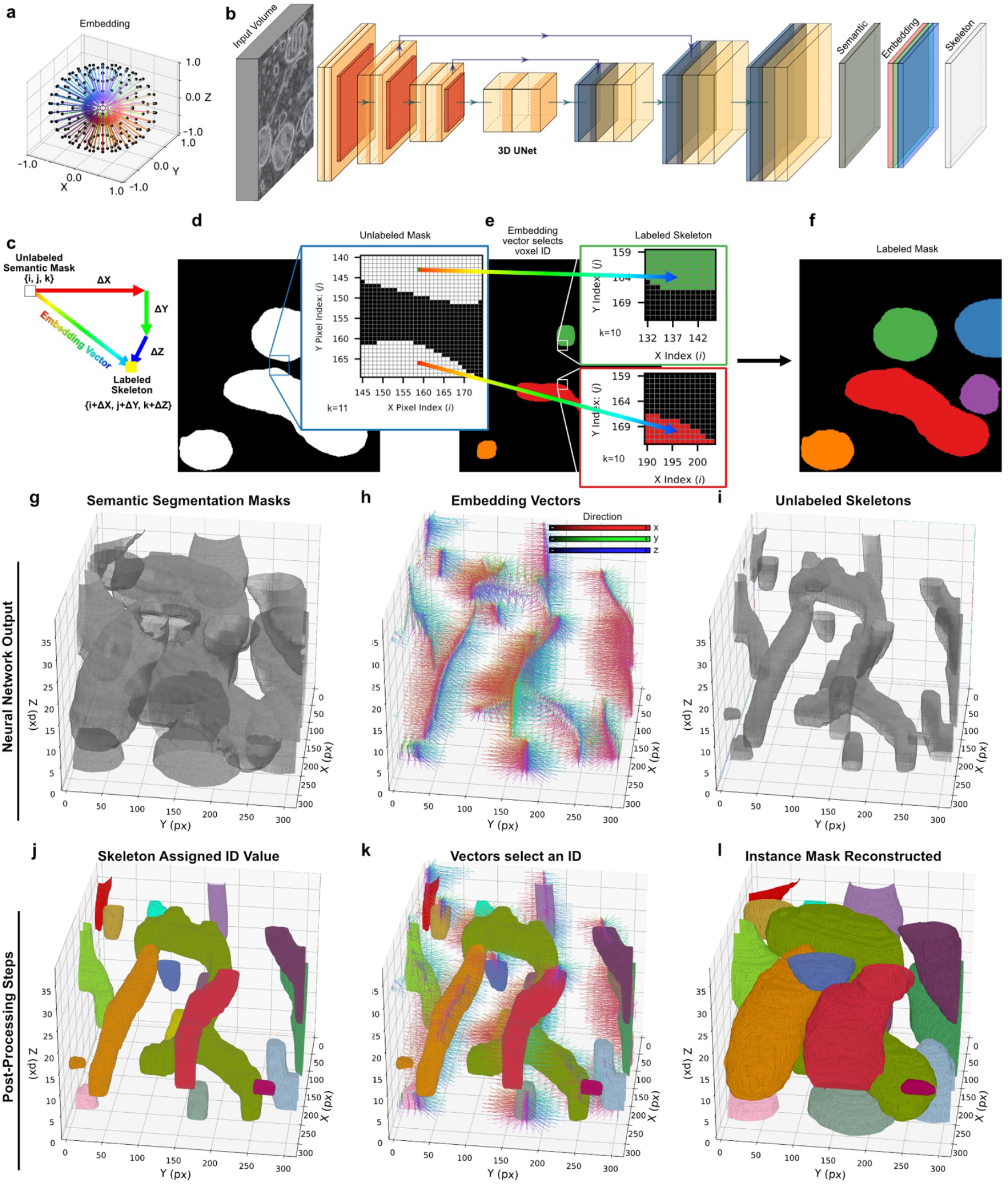
Overview of instance segmentation procedure via neural network skeleton embedding. (**a**) A schematic representation of spatial embedding. Voxels of a sphere in 3D space (black dots) are associated with an object by vectors (color-coded based on their orientation) pointing from their spatial location to the object’s local mean (grey). This principle is applied for mitochondria segmentation. (**b**) An input volume is passed through a trained 3D UNet neural network model that predicts a semantic segmentation mask, unlabeled skeletons, and 3D skeleton embedding vectors. (**c**) Each unlabeled voxel at location *ijk* with embedding vector *V*_*ijk*_ = [Δ*X*, Δ*Y*, Δ*Z*] points to a location *i* + Δ*X, j* + Δ*Y, k* + Δ*Z* which lies within the associated skeleton. (**d-f**) Simplified 2D representation of the model’s key steps. (**d**) Each voxel of the semantic mitochondria mask is assigned a 3D embedding vector predicted by the neural network, pointing to an unlabeled skeleton. (**e**) All predicted skeletons are then labeled using an efficient flood fill function. (**f**) Each voxel is assigned the ID of the skeleton it points to. (**g-i**) 3D representations of the model’s key steps. A pretrained deep-learning model directly infers the following, rendered in 3D for illustration purposes: (**g**) semantic segmentation mask, (**h**) 3D embedding vectors (color-coded for orientation), and (**i**) unlabeled skeletons. From these deep neural network outputs, the instance segmentation masks are generated. (**j**) skeletons are assigned unique IDs via flood fill. (**k**) The embedding vectors project onto labeled skeletons, assigning an ID to each voxel. (**l**) 3D rendering of fully segmented and labeled mitochondria instances.

When tasked with segmenting mitochondria in a 3D FIB-SEM dataset, SKOOTS approaches the speed of affinity-based segmentation while maintaining the boundary invariance characteristic of flow-based methods. Once skeletons are labeled with unique IDs, they must be associated with the voxels that make up the full object. SKOOTS operates by predicting a 3D optical flow - also referred to as spatial embedding vectors - using a deep neural network (**Figure 1a-c**) for each voxel in a semantic segmentation mask of mitochondria (**Figure 1d**). These vectors direct each voxel to a uniquely labeled, explicitly predicted skeleton (**Figure 1e**). Instead of computing these vectors by iteratively flowing up the gradient of an Eikonal distance function, as in *Omnipose*, we predict them directly. This contributes to SKOOTS’s increased segmentation speed and memory efficiency. Finally, voxels embedded within a labeled skeleton are assigned to that skeleton’s ID, and reversing the embedding yields a complete instance mask (**Figure 1f**).

The steps of SKOOTS can be summarized as follows. First, a pretrained neural network predicts three outputs: (1) a semantic segmentation mask of mitochondria (**Figure 1g**), (2) a set of 3D skeleton embedding vectors (**Figure 1h**), and (3) an unlabeled segmentation mask representing distinct mitochondria skeletons (**Figure 1i**). Embedding vectors are predicted for every voxel in the volume and represented as a 3D RGB image, where the red, green, and blue values reflect the embedding strength in the X, Y, and Z dimensions, respectively. Next, a flood fill algorithm is applied to the unlabeled skeletons to assign a unique ID to each instance (**Figure 1j**). Following this step, each voxel’s embedding vector points to a labeled skeleton with a unique ID (**Figure 1k**), and the voxel is assigned that ID. An instance mask is then reconstructed from these assignments (**Figure 1l**). For large datasets, this entire process can be parallelized, further improving efficiency.

Most datasets generated using 3D EM or fluorescence imaging are anisotropic: the pixel size in X-Y is often smaller than the Z step. SKOOTS predicts embedding vectors within a normalized range of −1 to 1 across all spatial dimensions. By scaling these vectors proportionally to the anisotropy of the dataset, SKOOTS can accurately segment both isotropic and anisotropic volumetric datasets generated using FIB-SEM, serial blockface SEM, confocal microscopy, or any other 3D imaging technique.

### A new annotated dataset for FIB-SEM mitochondria segmentation

A major limitation in developing accurate 3D segmentation approaches is the scarcity of open-access training datasets. While some high-quality 3D segmentation datasets are available^19^, their usefulness can be limited by differences in mitochondrial morphology across cell types, which are often highly heterogeneous.

We generated a high-resolution FIB-SEM dataset of early-postnatal murine cochlear sensory epithelium, containing sensory hair cells and Deiter’s cells, with a total volume of 23.4 µm (X) × 18.5 µm (Y) × 11.1 µm (Z). The voxel size was 3.82 nm in X and Y, with a 20 nm milling step in Z, resulting in an anisotropy ratio of approximately 1:5 and an overall image size of 6194 (X) × 4849 (Y) × 553 (Z) voxels. The mitochondria within this volume display a range of morphologies, sizes, and packing densities.

For training SKOOTS segmentation, and validation of performance, we manually annotated 722 mitochondria from a single outer hair cell within a subvolume measuring 1968 (X) × 3528 (Y) × 201 (Z) voxels. We employed a human-in-the-loop annotation paradigm, in which instance masks generated by a semi-supervised segmentation approach were iteratively corrected by expert annotators. Exemplar mitochondria are shown in **Figure 2**. The average volume of annotated mitochondria was 0.038 ± 0.023 µm^3^, with an average surface area of 0.691 ± 0.381 µm^2^. Special attention was given to accurately segmenting densely packed mitochondria, which are typically the most challenging to segment.

**Figure 2.**
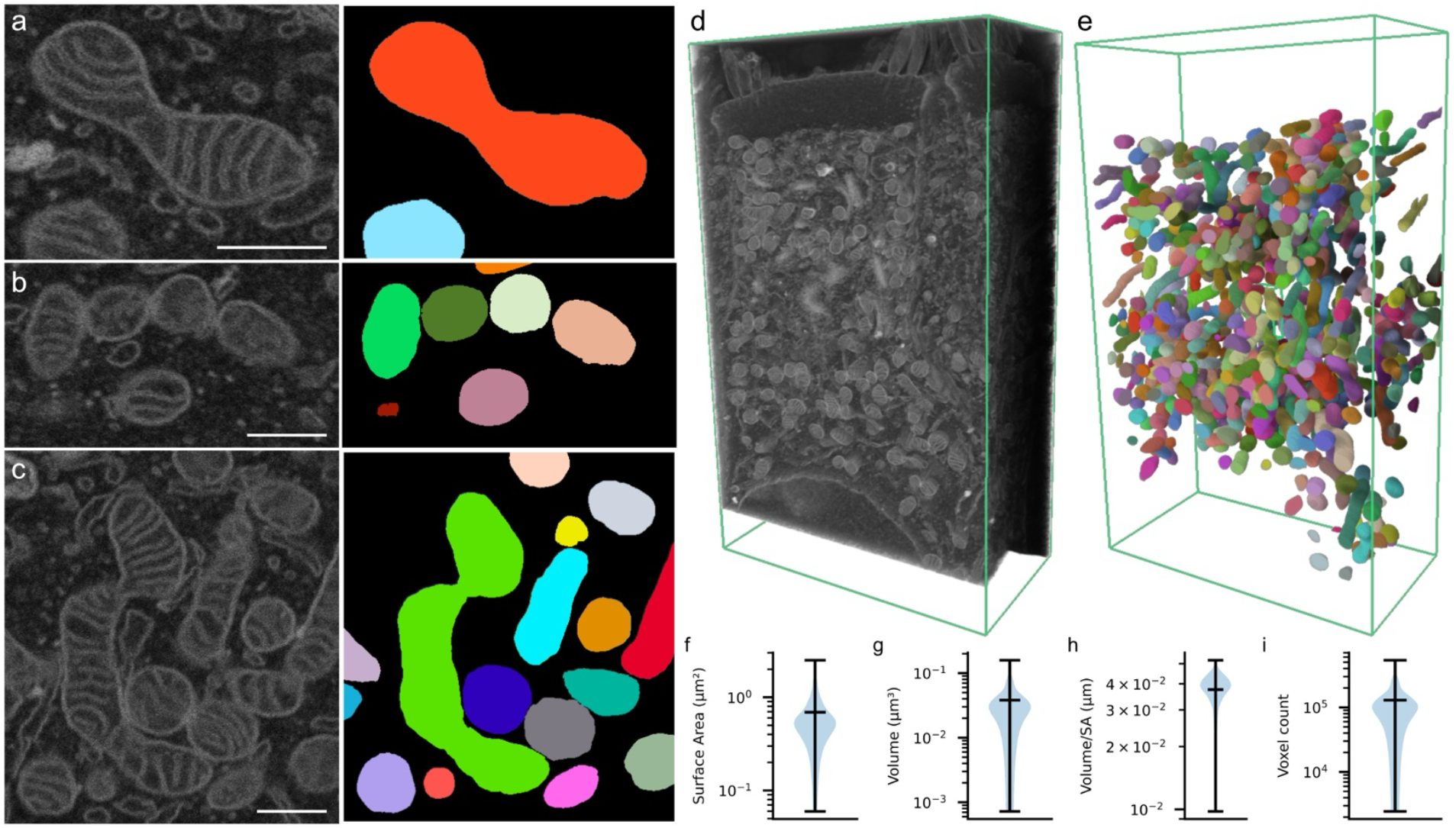
Manually annotated FIB-SEM mitochondria instance segmentation training dataset from an early postnatal murine outer hair cell. (**a**) Representative two-dimensional slice showing exemplar mitochondria (*left*) and corresponding segmentation mask (*right*), with distinct mitochondria represented by different colors. (**b**) Exemplar image and associated mask from a region containing densely packed mitochondria with challenging outer membrane boundaries. (**c**) Example of a mitochondrion (*lime green*) exhibiting atypical morphology. (**d**) Three-dimensional rendering of the full training dataset. (**e**) Three-dimensional rendering of approximately 700 manually annotated ground truth mitochondria instance masks. Using this dataset, we computed distributions for: (**f**) surface area, (**g**) volume, (**h**) volume-to-surface area ratio, and (**i**) total voxel count per mitochondrion. All scale bars represent 500 nm. Data are shown as Mean ± 15^th^ and 85^th^ percentile.

We divided the annotated dataset into training and validation sets along the Z-axis. The first 151 slices were used for training, and the remaining 50 slices for validation. The validation set contained 198 manually annotated mitochondria, accounting for 27% of all labeled instances and 24.8% of the total annotated volume.

### SKOOTS enables accurate 3D segmentation on a large scale in FIB-SEM volumes

Using our annotated mitochondria dataset and neural network architecture, we trained a deep learning model to perform instance segmentation with SKOOTS. When evaluated against a validation set of 198 manually annotated mitochondria, SKOOTS achieved high segmentation accuracy across a range of mitochondrial sizes and morphologies (**Figure 3a**). A representative 2D slice showing predicted skeletons and the resulting segmentation masks is presented in **Figure 3b-e**. While most mitochondria were segmented correctly (see 3D renderings in **Figure 3f-g**), occasional errors in skeleton prediction led to segmentation errors (**Figure 3h**). Despite these errors, the predicted segmentation masks preserved the overall distribution of mitochondrial morphological features, matching those derived from the manually annotated dataset (**Figure 3i, 3j**).

**Figure 3.**
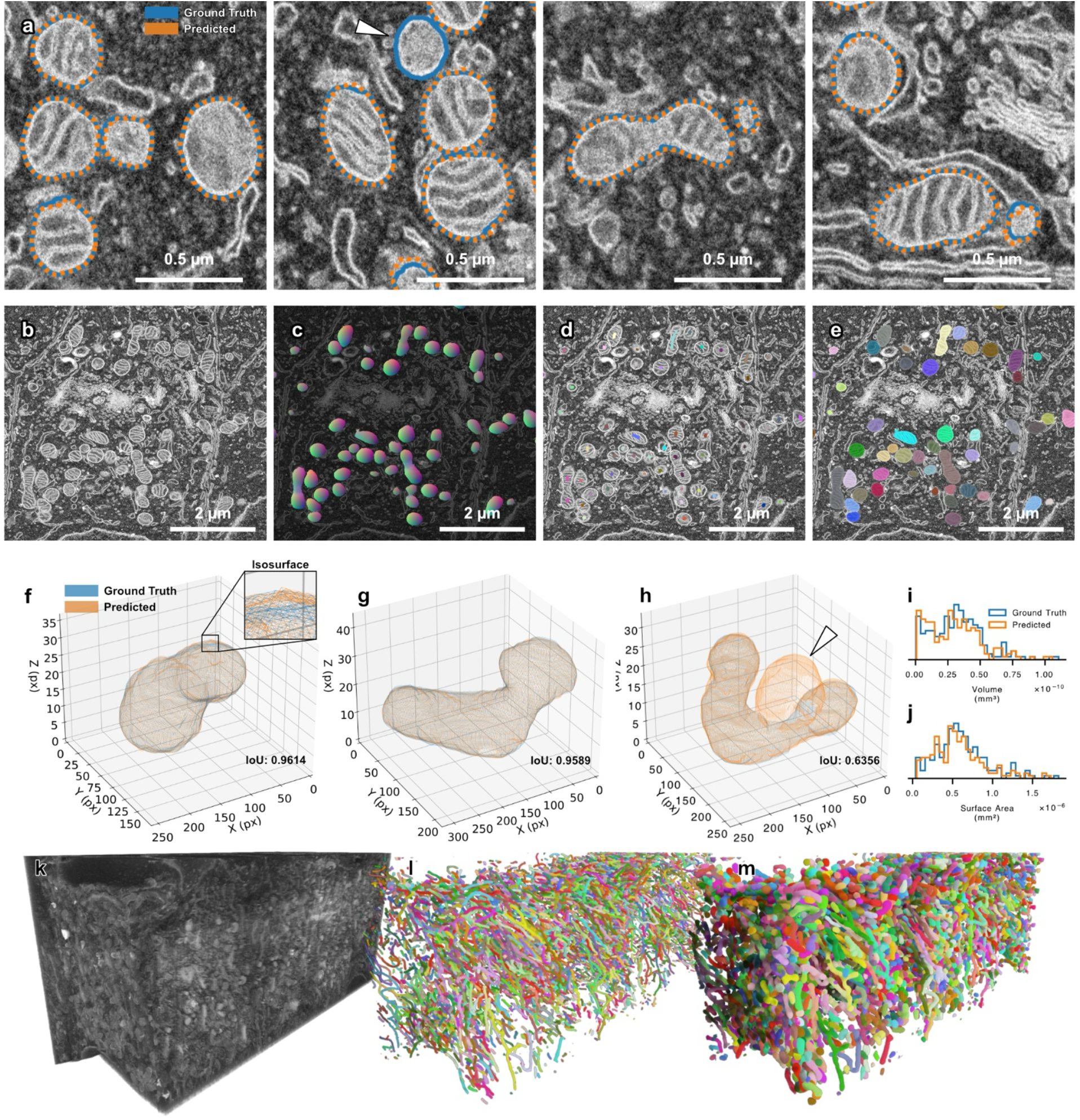
SKOOTS enables accurate instance segmentation of large-scale datasets. (**a**) Representative 2D slice showing a segmentation result. Predicted segmentation outlines (*orange dashed*) are overlayed on ground truth segmentation outlines (*blue*). A missed mitochondrion instance is indicated by a *white arrowhead*. (**b-e**) Visualization of SKOOTS workflow: (**b**) Single FIB-SEM image slice, (**c**) Corresponding slice showing 3D skeleton embedding vectors, (**d**) Labeled skeletons, (**e**) Resulting segmentation masks. Individual instance IDs are represented by different colors. (**f**) 3D mesh rendering comparing ground truth (*blue*) and predicted (*orange*) segmentation mask isosurfaces. (**g**) SKOOTS accurately segments mitochondria with diverse morphology. (**h**) Example of an under-segmentation error. (**i**) Histogram of mitochondria volumes calculated from manually annotated (*blue*) and predicted (*orange*) segmentation masks. (**j**) Histogram of mitochondria surface areas for the same sets. (**k**) 3D rendering of the FIB-SEM input volume. (**l**) 3D rendering of predicted skeleton instances. (**m**) 3D rendering of final mitochondria instance segmentation masks.

To demonstrate scalability, we applied SKOOTS to the full, multicellular FIB-SEM dataset (**Figure 3k-m**). On consumer-grade hardware, SKOOTS segmented 6,897 mitochondria within a 6194 (X) × 4849 (Y) × 553 (Z) voxel 3D EM dataset at 3.82 nm/px resolution (X and Y) in 25.8 minutes, averaging 267 segmented mitochondria per minute.

### SKOOTS enables more accurate mitochondria segmentation compared to other automated approaches

To evaluate the performance of SKOOTS, we trained an affinity-based segmentation model using the same neural network architecture and dataset, applying a shared set of parameters to ensure an unbiased comparison. We then applied this model to our manually annotated validation dataset, computed instance masks, and assessed segmentation performance **(Figure 4, 5)**. The affinity-based approach frequently misclassified contacting mitochondria as a single object due poor affinity graph predictions at membrane contact points, resulting in under-segmentation errors (**Figure 4c, 4d**). In contrast, SKOOTS (**Figure 4e-g**) predicted distinct skeletons for each mitochondrion, enabling better separation of touching mitochondria and resulting in fewer under segmentation errors. Over identical conditions for our validation dataset, affinity segmentation resulted in an under-segmentation rate of 1.17% vs 1.04% with SKOOTS (**Figure 5**).

**Figure 4.**
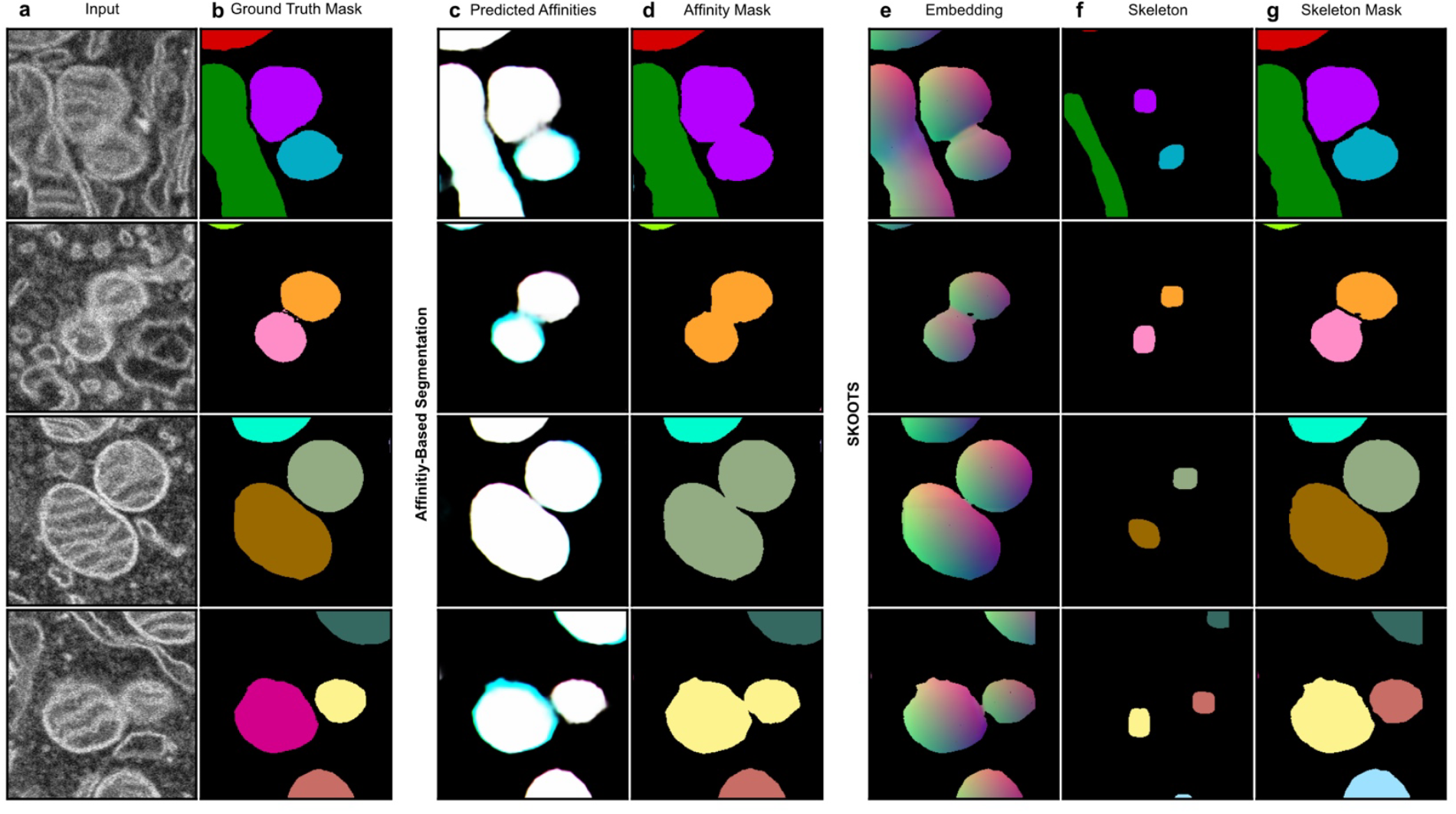
SKOOTS outperforms affinity-based segmentation in instances of challenging mitochondria boundaries. (**a**) Representative FIB-SEM images highlighting difficult segmentation cases, (**b**) Corresponding ground truth segmentation masks for each image. Images and masks were processed by two segmentation pipelines: (**c-d**) Affinity-based segmentation and (**e-g**) SKOOTS. (**c**) Predicted affinity graphs (**d**) Resulting instance masks; weak boundary predictions led to under-segmentation. (**e**) SKOOTS-predicted embedding vectors and (**f**) Predicted skeletons (**g**) Final segmentation masks. SKOOTS more accurately segments adjacent mitochondria, reducing under-segmentation errors.

**Figure 5.**
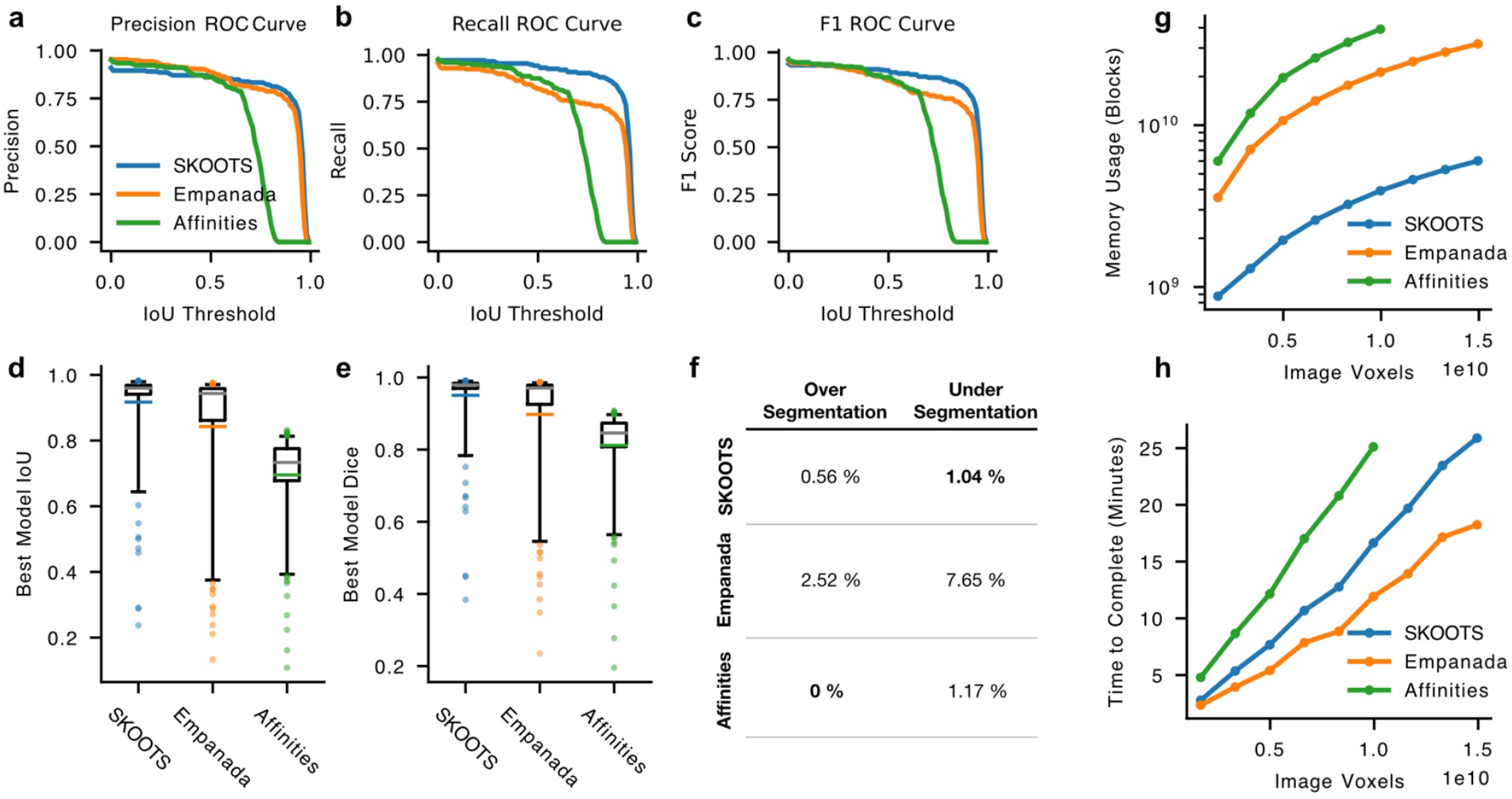
*SKOOTS* is accurate, fast, and memory-efficient at segmenting mitochondria compared to other segmentation approaches. Mitochondria in our validation FIB-SEM dataset were segmented using *SKOOTS* (*blue*), Empanada (*orange*), and an Affinity-based (*green*) segmentation approaches, and compared to manually annotated ground-truth segmentation masks for accuracy. Segmentation performance was assessed by calculating (**a**) Precision, (**b**) Recall, and (**c**) F1 score for each approach at multiple rejection thresholds to generate region operating characteristic (ROC) curves for each metric. The average intersection over union (IoU, **d**) and Dice (**e**) score were computed for all correctly identified mitochondria; colored horizontal lines represent mean values, while grey lines represent median values. (**f**) Over- and under-segmentation error counts for each approach, with best-performing values shown in bold. (**g**) Total GPU memory usage, and (**h**) execution time required to segment volumes of varying total size. All models were trained on the same dataset for equivalent durations. *SKOOTS* and Affinity-based segmentation used identical network architectures and model evaluation scripts. *Empanada* was trained and evaluated using its native Napari plugin.

We also fine-tuned Empanada^38^, a pseudo-3D mitochondria segmentation pipeline that employes 2D optical flow followed by 3D reconstruction, on our dataset and evaluated its performance on the same validation set. Unlike the affinity-based approach, Empanada uses a different network architecture and was pretrained on external datasets. For three methods—SKOOTS, affinities, and Empanada—we computed standard detection and segmentation accuracy metrics (**Figure 5**). SKOOTS achieved the highest precision, recall, and F1 score for mitochondria detection (**Figure 5a-c**), and outperformed the other methods in average intersection over union (IoU) and Dice coefficient metrics (**Figure 5d-e**), indicating superior segmentation accuracy. Additionally, SKOOTS had the lowest rate of under-segmentation errors (**Figure 5f**).

SKOOTS also required less GPU memory than either Empanada or the affinity-based approach (**Figure 5g**). While SKOOTS was faster than the affinity-based segmentation approach, it was slower than Empanada, taking 25.8 min to complete 1.49×10^10^ voxels in our validation dataset compared to 18.2 min for Empanada. (**Figure 5h**). This difference may be due to more efficient acceleration of 2D deep-learning operations compared to 3D.

### SKOOTS can be applied for efficient segmentation of fluorescence light microscopy datasets

Although SKOOTS was originally developed for mitochondria segmentation in FIB-SEM volumes, it is a general-purpose algorithm for high-resolution 3D segmentation and can be applied across diverse imaging modalities. To demonstrate its versatility, we trained and applied SKOOTS to segment a 1000(X) × 1000(Y) × 60(Z) voxel confocal microscopy Z-stack of neonatal auditory hair cells immunolabeled for the cytoplasmic hair cell marker MYO7A (**Figure 6a**). These cells form a densely packed epithelium in four rows and, when labeled cytoplasmically, often exhibit poorly defined borders. Despite this, SKOOTS successfully predicted accurate 3D segmentation masks.

**Figure 6.**
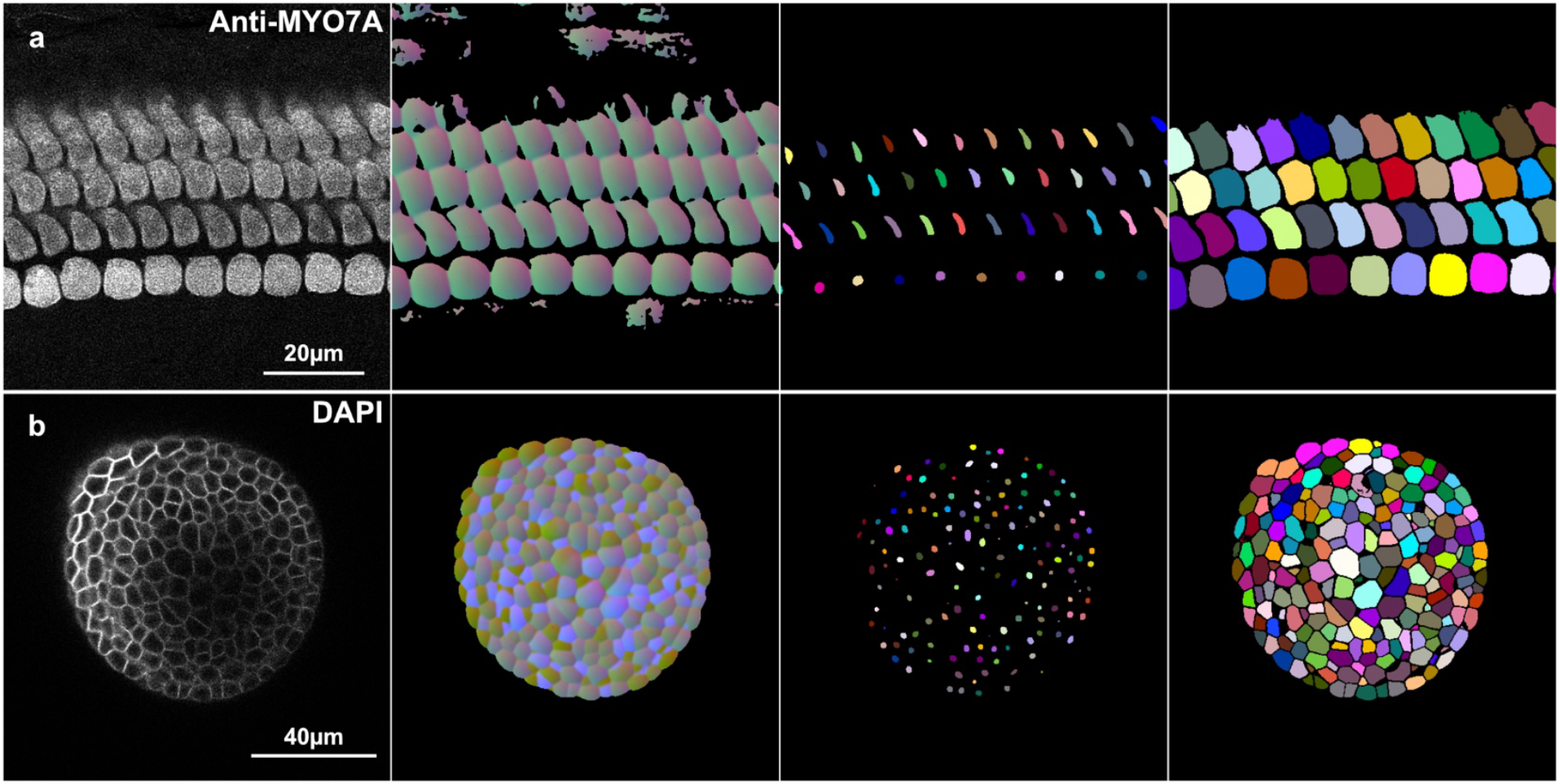
*SKOOTS* enables accurate segmentation of fluorescence microscopy datasets. *SKOOTS* is not limited to electron microscopy volumes and can be applied to a variety of 3D imaging modalities. We successfully trained and applied *SKOOTS* to segment: (**a**) auditory hair cells immunolabeled with anti-MYO7A, and (**b**) lateral root primordia cells of *Arabidopsis thaliana*, with cell walls stained with DAPI. Both samples were imaged using confocal microscopy. For each dataset, panels show (from *left* to *right*): the input image, predicted embedding vectors, predicted skeletons, and the resulting segmentation mask.

We further tested SKOOTS on an open access fluorescence microscopy dataset of Arabidopsis thaliana lateral root primordia, in which the cell walls are stained with DAPI, using a confocal dataset previously published and deposited by Adrian et al 2020^43^ (**Figure 6b**). Together, these results demonstrate that SKOOTS is a robust and efficient instance segmentation tool applicable to a range of three-dimensional imaging datasets.

### SKOOTS can identify morphological differences in mitochondria treated with aminoglycosides

To demonstrate the value of SKOOTS in biomedical research, we assessed its utility in identifying mitochondrial changes in cultured cochlear hair cells treated with aminoglycosides. Aminoglycosides are potent antibiotics, however, prolonged treatment with aminoglycosides can lead to permanent sensorineural hearing loss as a result of hair cell death. The ototoxic effect of aminoglycosides in hair cells is multifaceted, and include disrupting lipid membranes and increasing mitochondrial reactive oxidative species production^13,18,44,45^. We used this model to evaluate whether SKOOTS could reveal early morphological changes in mitochondria during aminoglycoside exposure.

In order to assay aminoglycoside induced hair cell death and evaluate changes in mitochondria morphology we incubated early postnatal cochlear cultures with 10 µM gentamicin *in vitro* for either 24 or 48 hours (**Figure 7**). As reported previously^46^, outer hair cell death was observed in the basal region of the cochlear after 48 hours of gentamicin treatment, compared to controls with no gentamicin. However, no hair cell loss was observed after 24 hours (**Figure 7a**). Cochleograms generated using the hair cell analysis toolbox (HCAT) ^47^, demonstrate the expected gradient in hair cell death from the base to apex of the cochlea after 48 hours treatment (**Figure 7b-c**). Visualization of outer hair cell mitochondria in the mid-basal region of the cochlea by TEM revealed no obvious morphological differences after 24-hour gentamicin treatment, with only cell debris following 48-hour gentamicin treatment (**Figure 7d**).

**Figure 7.**
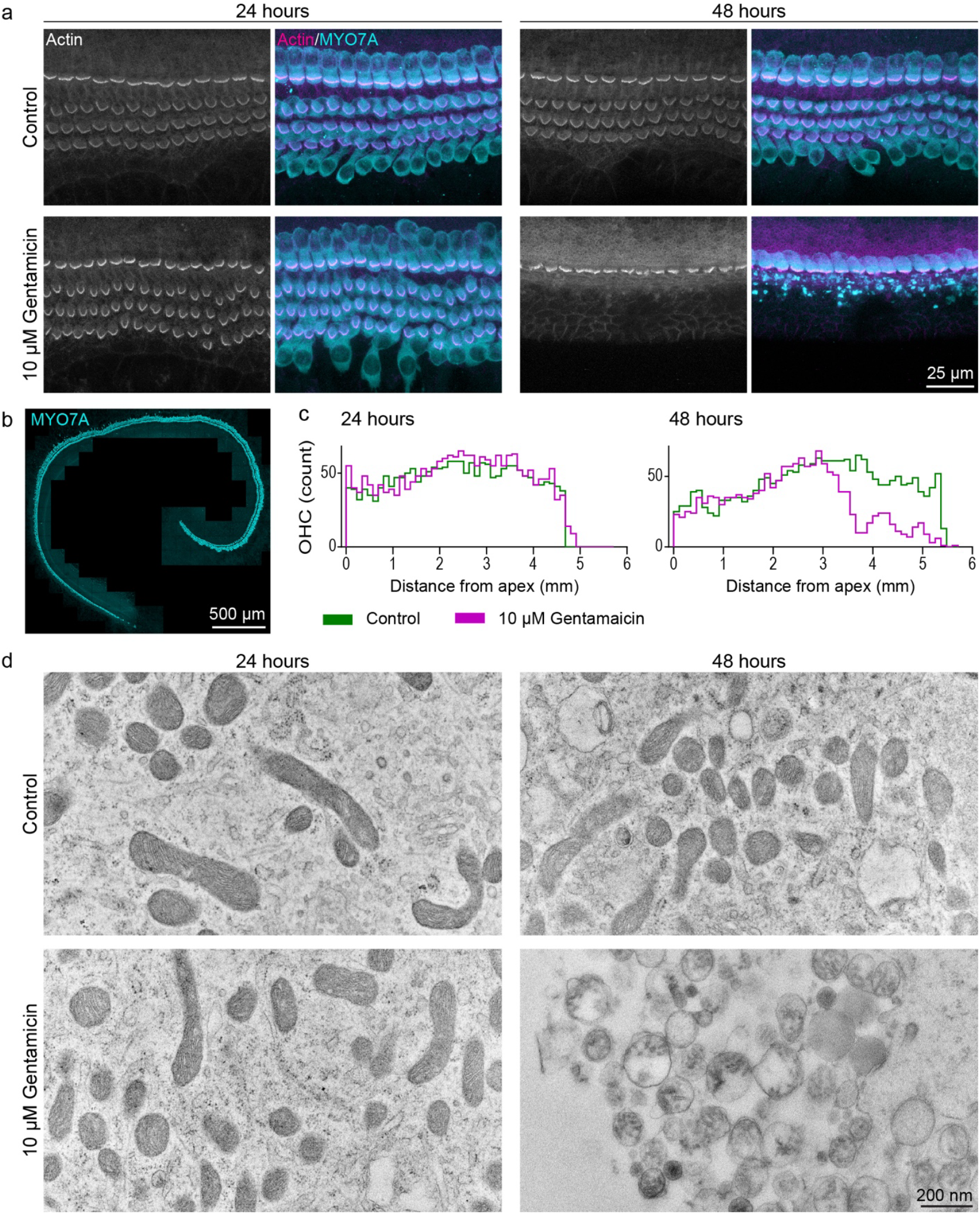
Classical TEM reveals no overt change to mitochondria following 24 hour gentamicin exposure. Postnatal day 2 mouse cochlear explants were cultured 1 day *in vitro*, then incubated with 10 µM gentamicin for either 24 or 48 hours. Control cultures were incubated without gentamicin. **a**, Confocal z-projections of hair cells from the mid-basal region of the cochlear coil reveal OHC loss after 48 hours of gentamicin treatment, while no loss is observed after 24 hours. *Scale bar*, 25 µm. **b**, Tiled confocal micrograph of a cochlea treated with gentamicin for 48 hours, showing the extent of hair cell loss. *Scale bar*, 500 µm. **c**, Quantification of OHCs using our previously reported deep learning cell detection model, HCAT, reveals a gradient of OHC loss from base to apex after 48 hours of treatment, with no detectable OHC loss at 24 hours **d**, TEM micrographs of OHCs from the mid-basal region reveal no overt morphological disruption of mitochondria at 24 hours of gentamicin treatment compared to untreated controls. After 48 hours, severe morphological disruption is observed in mitochondria within cellular debris. *Scale bar*, 200 nm. TEM images are representative of >3 OHCs per condition.

To assess for any subtle disruptions to mitochondria morphology, not apparent from TEM imaging, we collected FIB-SEM volumes of outer hair cells in the mid-basal region of cultures treated for 24 hours with gentamicin (5 cells), and control (3 cells). The resulting datasets were segmented using SKOOTS (**Figure 8**). We computed the volumes and surface areas of the resulting 3D instance masks using Dragonfly image analysis software^48^. To account for variability in treatment effects on cells within the same cochlea, we modeled mitochondrial volume and surface area using a Bayesian hierarchical framework (**Supplemental Figures S2-S8**), employing uninformative priors. A total of 8,728 mitochondria were included in the model to infer posterior predictive distributions. The analysis revealed a statistically significant effect of aminoglycoside treatment on both mitochondrial volume and surface area following gentamicin treatment.

**Figure 8.**
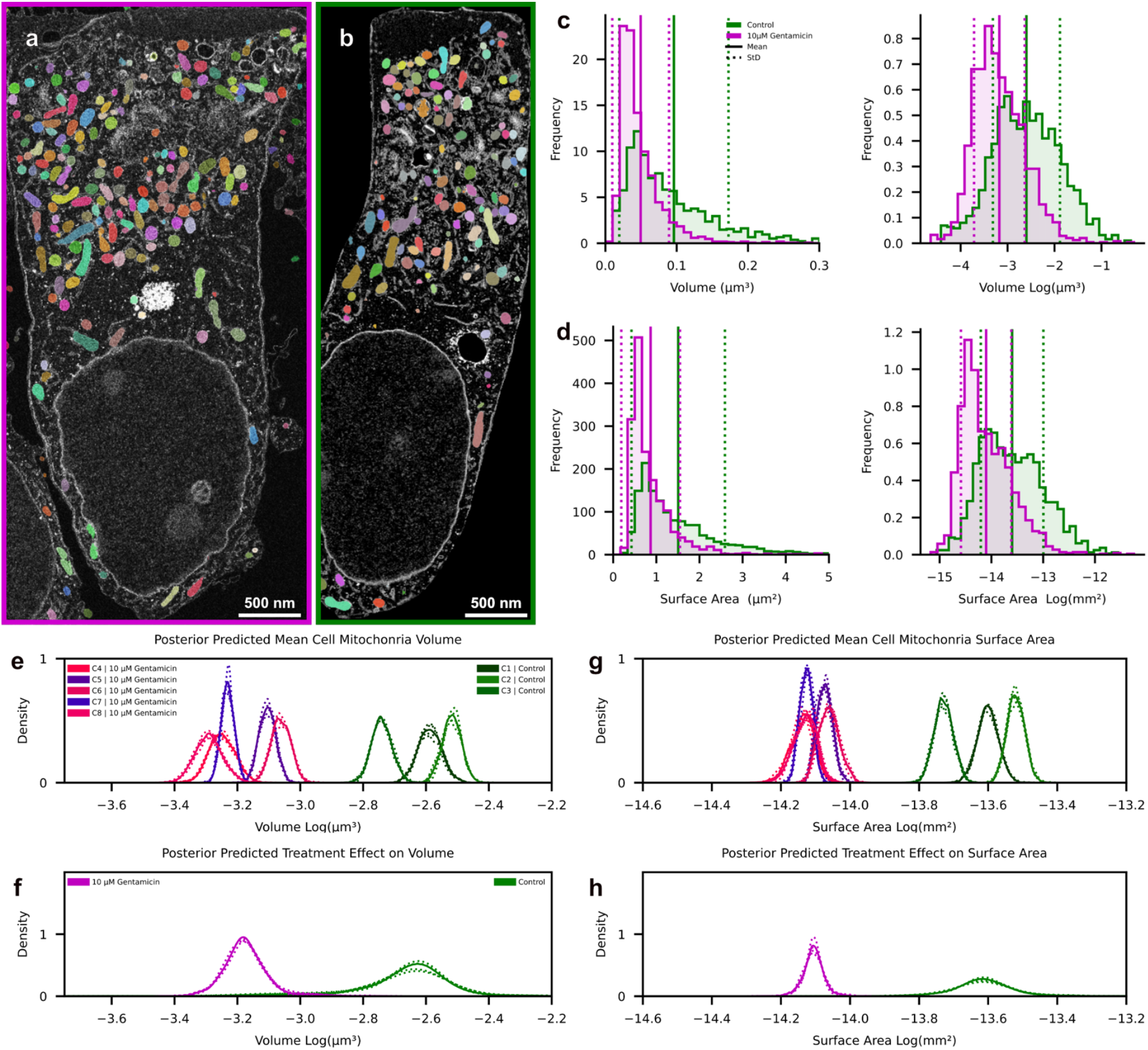
SKOOTS reveals morphological changes in outer hair cell mitochondria following gentamicin treatment. Three-dimensional mitochondrial masks were generated from FIB-SEM volumes of cochlear hair cells following SKOOTS segmentation. Five hair cells from a cochlea treated with 10 μM of Gentamicin (**a**) and three hair cells from an untreated control (**b**) were analyzed. **c**, Mitochondrial volume distribution show a leftward shift following gentamicin treatment (*left*), which is more apparent when plotted on a logarithmic scale (*right*). **d**, Mitochondrial surface area also decreases following treatment. A Bayesian hierarchical model was used to assess treatment effects while accounting for cell-to-cell variably. **e, g**, Posterior distributions of cell-specific mean mitochondrial volume (**e**) and surface area (**g**) reveal within cell variation. **f, h**, The model reveals statistically significant reduction of mitochondrial volume (**f**) and surface area (**h**) following gentamicin treatment.

Posterior predictive checks confirmed a good model fit to the observed data, and no divergences were encountered during sampling. Here, therefore, we demonstrate that analysis of FIB-SEM datasets by SKOOTS enables the identification of mitochondria disruption, that it was not previously feasible to obtain.

## DISCUSION

Here, we present SKOOTS—a 3D deep learning approach for mitochondria segmentation. By explicitly predicting object skeletons, SKOOTS achieves high accuracy, memory efficiency, and fast execution. It was developed to enable segmentation of large, densely packed objects that are challenging for existing 3D optical flow or boundary detection approaches. While originally designed for mitochondria segmentation in EM datasets, SKOOTS is generalizable across imaging modalities and object types, representing a significant advancement in 3D biomedical image segmentation. This work was motivated by our need to study mitochondrial population dynamics in auditory hair cells. These post-mitotic cells have exceptionally high energetic demands, leading to an abundance of densely packed mitochondria—conditions under which existing 3D EM segmentation approaches, both semi-automatic and automatic, perform poorly.

Modern graphics cards can rapidly evaluate neural networks; however, many segmentation approaches require additional algorithms to convert network outputs into instance segmentation masks. These post-processing steps often become the limiting factor for 3D segmentation speed. For example, *Cellpose* and *Omnipose* rely on clustering algorithms, most commonly DBSCAN, to assign unique ID labels. DBSCAN has a time complexity of O(N^2^)^37^, making it exponentially slower in three dimensions. Segmenting a 100(x) × 100(y) × 100(z) voxel image volume using DBSCAN is approximately 100,000 times slower than segmenting a single 100(x) × 100(y) pixel 2D image.

In contrast, boundary detection networks use watershed-based algorithms on predicted affinity graphs to assign ID labels. Affinity graphs represent connections between voxels within the same object; voxels belonging to different objects are not connected, which effectively creates sub-voxel boundaries between them. When all affinities are correctly predicted, these sub-voxel boundaries delineate objects with high precision. This watershed-based approach scales linearly with input size and is therefore more efficient for 3D processing^49^. However, even small errors in the affinity graph, such as a gap between two boundaries, can significantly degrade performance, leading to under-segmentation and poor segmentation performance. Manual correction of watershed segmentations following semantic mask prediction improved accuracy, and is faster than full manual annotation, but remains impractical at scale.

SKOOTS does not seek to supersede other segmentation algorithms in all use cases. Rather, we offer superior performance in a context poorly served by either: segmentation of high-resolution, densely packed objects with poorly delineated object boundaries. When comparing the performance of SCOOTS to existing methodologies, a neural network’s accuracy depends on multiple factors beyond algorithm design, including network architecture, training data, and post-processing. To better isolate the contribution of the segmentation strategy itself, we controlled for these variables where possible in our comparisons. Nonetheless, without fully re-implementing other pipelines, subtle approach-specific optimizations may still account for some observed differences.

SKOOTS excels in high-resolution instance segmentation when object diameters are large enough to support the prediction of well-defined skeletons. Consequently, there is a practical lower limit to object diameter in pixels: if objects are too small, skeletons may merge or fragment, causing under-or over-segmentation, respectively. In practice, this issue does not arise when imaging at sufficiently high resolution. Although the core operations in SKOOTS are dimension-agnostic and can be applied in 2D, the computational advantage of using flood fill (as in SKOOTS) over DBSCAN (as in *Omnipose*) becomes negligible in 2D. Thus, SKOOTS offers no clear benefit over *Omnipose* for 2D segmentation tasks. In 3D, however, SKOOTS requires significantly less GPU memory than *Omnipose*. However, SKOOTS encodes four values per voxel (embedding vectors in X, Y, Z and a skeleton mask), compared to three for affinity-based methods (affinities in X, Y and Z). This results in a 33% increase in memory usage. For our dataset—32.6 GB at native resolution and bit-depth—this increase was acceptable. For substantially larger datasets, however, this may present a bottleneck unless mitigated by additional features, such as disk-based data streaming.

While we implemented SKOOTS in *Python* due to its flexibility and the mature deep learning ecosystem, *Python* is not optimized for speed. In our segmentation pipeline, the majority of computational time is spent not on deep learning inference, but on conventional post-processing algorithms. Many of these operations could be significantly accelerated if reimplemented in compiled languages such as *C++* or *Rust*. A dedicated implementation could therefore yield substantial speed gains over our native *Python* code and further enhance usability.

The original FIB-SEM dataset we generated was sufficient to train a model with high segmentation accuracy for our dataset. However, segmentation performance may degrade with changes in sample preparation, imaging conditions, resolution, or signal-to-noise ratio. Mitochondria also exhibit diverse morphologies across different tissues and cell types, and our dataset, derived from cochlear hair cells, may not fully represent this diversity. Notably, our training dataset lacks mitochondria in obvious pathological states such as swelling, vacuolization, or cristae degradation. As a result, the current model may be limited in its ability to segment or recognize abnormal mitochondria without further training. While classification-based approaches using deep learning could potentially identify mitochondrial morphology without requiring full segmentation, such methods are inadequate for extracting precise morphological statistics ^50^.

We have made the SKOOTS source code, neural network model, and all pretrained models publicly available. The code is implemented as a flexible *Python* library that can be easily integrated into existing segmentation pipelines. In addition, we provide comprehensive documentation detailing model training, evaluation, and code functionality. Given the inherent variability in sample preparation, imaging modality, anisotropy, and resolution across datasets, users will likely need to fine-tune or retrain their own models using their own annotated data. While manual annotation can be labor intensive, our pretrained models offer a strong starting point and significantly reduce the amount of data needed for training models that achieve high segmentation accuracy. Step-by-step training scripts and detailed guidelines are included to support external users.

In summary, SKOOTS bridges the gap between segmentation performance and accuracy in challenging 3D imaging conditions, delivering fast, memory-efficient, and highly accurate results. By advancing beyond prior optical flow techniques through direct skeleton prediction, SKOOTS offers a robust, generalizable framework for high-resolution 3D instance segmentation across a range of imaging modalities.

## MATERIALS AND METHODS

### FIB-SEM Data Collection

The original FIB-SEM dataset used in this study for model training and validation was collected as part of a separate study, detailed fully elsewhere^51^. In summary, early postnatal mouse organotypic cochlea cultures were utilized for an aminoglycoside damage repair assay, prior to fixation, processing, and imaging by FIB-SEM. Briefly, cultures were prepared from wild-type CD-1 mice at postnatal-day 2 (P2) as described previously^52^ and maintained *in vitro* for 1 day. Following a 15-minute exposure to 1 mM neomycin in HEPES-buffered HBSS (HBHBSS), cultures were washed with cold HBHBSS then incubated on ice at 4^°^C in HBHBSS with 0.5 mg/ml cationized ferritin for 15 minutes to track endocytosis. Cultures were washed with HBHBSS and allowed to recover in culture medium at 37^°^C/5% CO_2_ for 15 minutes, prior to fixation. It is not anticipated that this assay will disrupt mitochondria imaging or analysis. Animal work was carried out at the University of Sussex in full accordance with United Kingdom Home Office regulations.

Cultures were fixed for 2 hours at room temperature with 2.5% glutaraldehyde, 2 mM calcium chloride in 0.1 M sodium cacodylate buffer; followed by secondary fix with 1% osmium tetroxide, 1.5% potassium ferrocyanide in 0.1 M sodium cacodylate buffer for a further 2 hours. Samples were dehydrated by ethanol series and transferred to propylene oxide prior to embedding in an Epoxy Embed812/Araldite 502 resin mix.

Following tissue exposure by ultramicrotomy, FIB-SEM was carried out at the Center for Nanoscale Systems, Harvard University, on a FEI Helios Nano lab 660 operated at 3 kV. Milling and imaging was carried out in an orientation along the apical/basal axis of the organ of Corti, orthogonal to the reticular lamina, travelling through multiple outer hair cells in row one simultaneously. Three full outer hair cells in row one in the mid region of the coil were imaged in a serial FIB-SEM volume, 23.4(x) 18.5(y) 11.1(z) µm, collected at 3.82 nm/px with 20 nm milling steps. The resulting image stacks were aligned in Dragonfly Software, Object Research Systems (ORS) Inc, Montreal, Canada, 2022.

### Gentamicin treatment assay, sample preparation, and imaging

The cochlear culture gentamicin treatment assay carried out to demonstrate the utility of SKOOTS is based on that published previously^46^. Briefly, postnatal day 2 cochlear cultures were prepared from wild type CD-1 mice, and maintained overnight in DMEM/F12 supplemented with 7% FBS and 10 ug/mL ampicillin. At P2+1 cultures were transferred into DMEM/F12 supplemented with 1.4% FBS, 2 ug/mL ampicillin and incubated with 10 µM gentamicin (Sigma G3632) or HBSS control for 24 or 48 hours at 37^°^C/5%. All procedures and protocols were approved by the Institutional Animal Care and Use Committee of Mass Eye and Ear and we have complied with all relevant ethical regulations for animal use.

Cultures visualized by light microscopy were washed with HBSS and fixed with 4% PFA in PBS for 1 hour. To visualize the hair cells, cultures were labeled with anti-Myosin 7A antibody (#25–6790 Proteus Biosciences, 1:400) and phalloidin-488 (Biotum CF488A, 1:100) overnight in PBS supplemented with 10% donkey serum, 0.1% triton X-100. Cultures were incubated for 4 hours with CF568 F(ab’)2 fragment of goat anti-rabbit (Biotum, 1:500) in PBS supplemented with 10% donkey serum, 0.1% triton X-100, prior to mounting in Prolong gold. Imaging was carried out on a Leica SP8 confocal microscope using a 63×1.3NA objective lens, either in single regions at high resolution or full coil images were collected using the tiling functionality of the Leica LASX acquisition software. Images were processed in Image J and Adobe CC. Hair cell counts and cochleogram generation was carried out using HCAT^47^.

Cultures processed for TEM and FIB-SEM underwent the same sample preparation as outlined for the initial FIB-SEM dataset, with the omission of potassium ferrocyanide from the secondary fix of cultures prepared for TEM. For TEM, ultra-thin (∼80 nm) sections were collected from the mid-basal region of the cochlear coils onto formvar/carbon coated slot grids. In order to visualize multiple OHCs from a single block >10 μm of tissue was removed from the block face by ultramicrotomy between rounds of ultra-thin sectioning. Sections were stained on grids with 1% uranyl acetate and lead citrate prior to imaging on a Hitatchi TH7800 TEM operated at 100 kV at the Center for Nanoscale Systems, Harvard University. Images were processed in Adobe CC.

For FIB-SEM, resin blocks were faced to exposure the tissue surface by ultramicrotomy. As previously, FIB-SEM was carried on a FEI Helios Nano lab 660 operated at 3 kV. EM-volumes were collected from the mid-basal region of the cochlear coils, collected at 3.82 nm/px with 20 nm milling steps. The resulting image stacks were aligned in Dragonfly Software, prior to mitochondria analysis by SKOOTS.

### Mitochondria dataset annotation

The mitochondria training and validation dataset consists of an 1968(x) 3528(y) 201(z) px excerpt of the larger FIB-SEM hair cell dataset. Data were annotated using Dragonfly Software, Object Research Systems (ORS) Inc, Montreal, Canada, 2022, through a “human in the loop” approach by predicting semantic segmentation masks, assigning labels via watershed, and manually correcting the results.

In brief, using the Dragonfly segmentation wizard, 3D semantic segmentation masks of mitochondria were manually generated for a small part of the training and validation data subset and used to train the in software neural network model which generated masks for the entire subset volume, separating each voxel into two labels: foreground (mitochondria) and background. A distance map was then generated from the semantic mask. Thresholds of the distance map served as seed pixels, which were then manually eroded to reduce over-segmentation. From these seeds, the watershed algorithm was used to fill the distance map basins and generate instance maps of mitochondria. While this approach was accurate for isolated, large mitochondria, the outcome was not reliable enough to serve as training data. When using this method, densely packed mitochondria tend to be under-segmented with poor borders, while small mitochondria were entirely missed. Therefore, the resulting masks were visually inspected for errors, and manually corrected.

### Ground truth skeleton calculation for training

To predict skeletons of objects using a neural network, we first needed to explicitly calculate ground truth skeletons for each manually annotated instance. Prior to skeletonization, we performed a linear interpolation on the binary mask of each image, scaling the instance closer to an isotropic pixel size. The skeleton was a binary image, where each pixel represents the local mean of the object. We applied scikit-image’s skeletonize function to each scaled instance of our annotated dataset. The nonzero values of each instance skeleton were saved as a file for later training. In the case of small instances where skeletonization fails, we substituted the skeleton for the object’s local center of mass. We found the quality of calculated skeletons affected the accuracy of segmentation, and therefore up/down-scaling the instance mask of an object prior to skeletonization could produce skeletons which are better defined and lead to improved segmentation.

### Data pre-processing and augmentation for training

Our training dataset consisted of multiple large, manually annotated images, their associated ground truth labeled mitochondria instance masks, and precomputed skeletons. These images were too large to pass through a deep learning model at once. Rather, crops of 300 × 300 × 20 voxels were randomly centered at an instance of mitochondria. To improve generalizability of our model, we applied a suite of augmentations to these crops which are broadly separated into morphological augmentations (rotation, shearing, and flipping) and image augmentations (brightness and contrast adjustment, addition of random noise, and randomly inverting the image), these were applied were applied randomly to the images, masks, and skeletons.

To reduce false positive predictions, we also explicitly included images in the training dataset which do not contain any instances. Rather, we chose areas which contained objects visually like mitochondria, such as vacuoles or other organelles, improving the model’s ability to discriminate such objects from mitochondria.

### Model architecture

As SKOOTS requires embedding vectors and skeletons to be well defined for segmentation accuracy to be high, implemented an efficient neural network architecture based on the ConvNext neural network architecture, which achieved performance superior to transformer-based models by applying a system of best practices to convolutional neural network design^40^. We applied the same techniques to the standard UNet architecture (**figure S1a-e**), trained, and compared the resulting model against other published architectures for biomedical image segmentation. We trained each model on a semantic segmentation task (for speed), varying the number of parameters (and therefore FLOPS), and compared accuracy (**Supplemental Figure S1f**,**g**), and graphic memory requirements (**Supplemental Figure S1h**).

The specific changes made were as follows: 1) we replaced each activation with a gaussian error linear unit^53^ (GeLU), increased each kernel size of each convolution from three to seven, 3) replaced each double convolution computational block with 3D ConvNext block (an inverted bottleneck design), and 4) substituted each batch normalization layer with layer normalization^54^. While the original ConvNext used a layer scale initialization factor of 0.1, we elected to use a value of 1 as we observed no increase in final performance yet a considerable increase in the number of epochs to reach a training loss plateau. Through these modifications, we achieved superior segmentation performance compared to competing segmentation architectures.

Our implementation allowed for dynamic changes in the number of computational blocks, number of dimensions at each block, amount of downscaling, and offers 2D and 3D implementations. Each model used to evaluate any segmentation approach had an identical structure with two downscale steps, two upscale steps, and two computational blocks at each step. Each of the previously reported architectures used in this study for comparison (UNet, UNet++, Cellpose Net) were re-implemented by convention, but allowed for dynamic adjustment in scaling model size. The source code and implementation details can be found here: https://github.com/indzhykulian/bism.

### Training procedure

Training was distributed over two GPU’s using the pytorch DistributedDataParallel model. Each training data image was cropped and augmented in a process outlined above and collated into a batch size of two crops of training data per GPU. We defined a single iteration over our dataset as 32 crops for each fully annotated training image. We trained the model for 10000 epochs with the AdamW optimizer with a weight decay of 1e-6 and a learning rate of 5e-4 which decays on a cosine annealing schedule with a period of 10000. The loss value for each output of the segmentation model, the semantic masks, skeletons, and embeddings, were calculated using the Tversky loss function, as this allowed for fine tuning the network’s sensitivity to false positive and false negative predictions.

Rather than directly pre-computing optimal vector fields and computing a loss value against these as in Cellpose and Omnipose, we instead calculated a loss value for each vector based on the distance of the resulting embedding from an optimal skeleton location. This allowed for more flexibility in the model predictions and avoids having to re-compute flow fields after each augmentation, which is incredibly costly in 3D. The general steps were as follows: 1) for each ground truth mitochondria voxel we calculated the closest skeleton, 2) after applying the vectors we calculated a gaussian loss based on the distance and a standard deviation factor. Formally, the embedding loss Φat location *ijk* with embedding *E*_*ijk*_ and its associated skeleton *S*_*ijk*_ is the following:

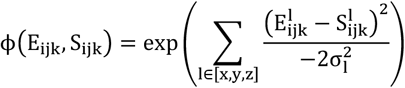

When expanded to three dimensions:

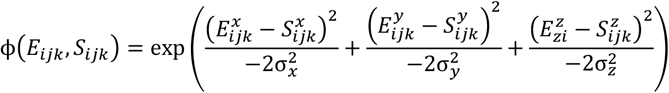

By applying this to each voxel we generated a probability map for each voxel, which we then compared to the ground truth semantic segmentation mask via the Tversky loss.

### Segmentation mask inference

SKOOTS applies the following steps for inference on new images. Arbitrarily large images are broken down into crops of identical size as training, which are then evaluated by a pretrained deep neural network model. The model predicts 5-channels of information: one channel for the semantic segmentation mask, one for the unlabeled skeleton, and three for the embedding vectors. Each embedding vector lies between −1 and 1, and therefore must be scaled such that any vector may be sufficiently large to point to a skeleton. Semantic segmentation masks are threshold at 0.5 and mask both the skeleton and embedding vectors, improving segmentation accuracy at the edges of objects. Vectors are scaled and stored, along with unlabeled skeleton masks, in pre-allocated arrays.

Following complete evaluation of the neural-network model, skeletons are assigned labels via flood fill. For speed, this process is also performed in crops, leading skeletons at the edges of a crop assigned multiple ID values, which are then detected and replaced such that each skeleton has one ID. This replacement occurs in place, avoiding copies and lowering the resulting memory footprint. Once each skeleton is assigned an ID, vectors select a voxels ID value by indexing the labeled skeleton mask. Formally, a neural network predicts 3 matrices: an unlabeled 3D semantic mask M, labeled skeleton mask *S* and predicted embedding vector *V* which all have shape (*X*_*in*_, *Y*_*in*_, *Z*_*in*_) where (*X*_*in*_, *Y*_*in*_, *Z*_*in*_) is the shape of the input image. Each value of these matrices may be indexed by the indices *ijk* where *i* is the X index and lies between 0 and *X*_*in*_, *j* is the Y index and lies between 0 and *Y*_*in*_ and k is the Z index and lies within 0 and *Z*_*in*_.

Therefore,

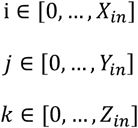

The value at each location of the embedding matrix V_*ijk*_ is defined as a vector which points to the closest voxel of a predicted instance skeleton. Therefore,

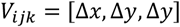

We may use these embedding vectors to assign labels to the unlabeled semantic mask if the probability at pixel location *ijk* is above some threshold *T*_*mito*_ (i.e., it belongs to a mitochondria). Formally,

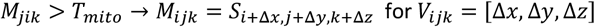

Simply, for each voxel of the input image, if that voxel represents a mitochondrion, there is an embedding vector which points it’s associated skeleton. As these skeletons already have a label, the complete instance mask of each mitochondrion can be reconstructed by selecting and ID at each location via the embedding vectors.

### Anisotropy

Anisotropic datasets occur when the absolute distance of a voxel a voxel represents is different along spatial dimensions. This commonly occurs when the resolution in Z of a volumetric imaging technique is lower than in X and Y. SKOOTS adapted to anisotropic data in two ways. First in training: Embedding vectors were predicted to lie at the closest skeleton. This was calculated as the minimum Euclidean distance between the location of the origin voxel, and all skeleton voxels. In some cases, this distance may force adjacent vectors to point in extremely different directions, usually in the Z direction. To combat this, the Euclidean distance from each voxel to the skeleton was scaled based on the anisotropy of the dataset, forcing the model to learn smooth embedding vector fields. Secondly, anisotropy was corrected at evaluation by scaling embedding vectors. Embedding vectors, when predicted, lie between −1 and 1. In a heavily anisotropic dataset, a pixel may need to be pushed in X and Y for tens of voxels to lie within a skeleton, but may be pushed in Z by one or two. With equal scaling of X, Y and Z, embedding vectors in the Z direction tended to be small and poorly predicted by the model, limiting overall performance. Therefore, we scaled embedding vectors such that all embedding vector predictions have similar minimum and maximum. In the case of our mitochondria dataset, we scaled the embedding vectors in X and Y by 60, and Z by 12. Formally, each embedding vector 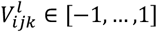 at location *ijk* and dimension *l* ∈ [*X, Y, Z*] is scaled by a scaling factor α^*l*^. Therefore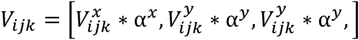.

### Segmentation of confocal images

Cochlear hair cell tissue was prepared as described in Buswinka *et al*. 2023^47^, and imaged with a Leica SP8 confocal microscope (Leica Microsystems) using a 63×, 1.3 NA objective. Confocal Z-stacks of 512 × 512 pixel images with an effective pixel size of 144 nm were collected using the tiling functionality of the Leica LASX acquisition software. All experiments were carried out in compliance with ethical regulations and approved by the Animal Care Committee of Massachusetts Eye and Ear. The dataset containing Arabidopsis thaliana lateral root primordia cell walls stained with DAPI was previously reported by Wolny et al. 2020^43^ and publicly available for download. Confocal datasets were manually annotated (when necessary) to generate training data and trained in an identical manner as above. As these datasets have differing pixel spacing, and therefore anisotropy, we adjusted the maximum embedding vector scaling accordingly. The brightness and contrast of the input images were adjusted prior to segmentation.

### Accuracy and morphology metrics

The accuracy of our segmentation approach was validated in two ways: 1) by the calculation of predicted instance intersection over union with associated ground truth segmentation masks, and 2) by the comparison of calculated morphology statistics. The intersection over union for each ground truth instance mask and associated predicted segmentation mask was calculated in *PyTorch*. The population morphology statistics were calculated in the Dragonfly software for both ground truth segmentation masks and predicted segmentation masks and further analyzed in python.

### Segmentation deep-learning architecture comparison

To select for the optimal architecture (the connections and operations of a deep-learning model) for our segmentation tasks, we validated our *UNet* + *ConvNext* blocks against other published architectures for biomedical segmentation, selecting the model with the highest performance for the least number of floating-point operations (FLOPS). Rather than training each model architecture on each respective segmentation task, we instead trained each on a semantic segmentation task, which is easier for a model to perform, and thereby trains faster. Each segmentation model architecture was implemented in *PyTorch* (see **Supplemental Methods**). Each model was trained with identical training parameters on our manually annotated mitochondria dataset. To use our data for a semantic task, we treated each uniquely labeled voxel as foreground, and each unlabeled as background. To facilitate reproducibility, we trained with identical random number generator seeding for each. To study the effect of model size, we started with a small number of convolution channels, and following training, doubled the channel count at each stage of the model and retrained.

This process continued until a model’s total memory requirements exceeded the avavilable video memory of our graphics cards (2×48 GB). FLOPS were calculated through the facebook research core vision library (fvcore, https://github.com/facebookresearch/fvcore) by passing a 300(x) 300(y) 20(z) voxel volume of random data through an untrained model. Model peak memory usage was calculated through *PyTroch’s* CUDA interface.

### Segmentation approach validation

Each segmentation approach may be evaluated on any deep neural network architecture, which affects accuracy and performance. Furthermore, training procedure, augmentations, hyperparameters, and other subtle features can affect overall approach accuracy. Therefore, to directly compare approaches, each were re-implemented to allow for identical training procedure and as similar inference as possible. Each aspect of training was defined by a single configuration file, with no more than necessary differences in training. Data, epochs, learning rate, data augmentations, model size, among other parameters, were kept identical for each approach. Implementation, along with training configuration files of may be found here: https://github.com/indhzkyulianlab/bism.

### 3D rendering

Two styles of 3D renders are presented in this work. **Figure 1g-I** and **figure 4f-h** were generated in matplotlib^55^ with a custom script. Briefly object meshes were approximated from input volumes (either segmentation masks or skeletons) by a marching cubes algorithm from scikit-image^56^. For speed, we used a step size of 2 and appropriate spacing to account for anisotropy. Each ID value was assigned a unique color and rendered in matplotlib. When appropriate, skeleton embedding vectors were rendered with the 3D quiver plot in Matplotlib, assigning a color based on the direction of the vector. For speed and clarity, only every 10^th^ vector was rendered in X and Y. Movies showing the rotations of these were generated by iterating the view angle of the 3D plot from 0 to 360, saving an image at each frame, then creating an animated GIF image in ImageJ^57^.

Renderings in **figure 3d, 3e** and **figure 4k-m** were generated by the Dragonfly Software. Multi-stack tifs of the input image, skeletons, and instance masks were imported into Dragonfly. A multi-ROI was generated from each greyscale ID value, and a random color was automatically assigned. The brightness and contrast of the image volume were adjusted. The multi-ROI of the skeletons and instance masks were overlayed on the input stack and rendered with Dragonfly’s 3D viewer.

### Statistics and Data Analysis

Mitochondria morphological descriptors were generated in the dragonfly software from masks predicted with our instance segmentation approach. Mitochondria volumes and surface areas were subdivided by treatment and cell ID. Bayesian models were constructed and fit with pymc, a python package for probabilistic modeling^58^. Each model was sampled using 4 chains for a total of 500 samples. *RHat* value, comparing the variance within each chain to the variance between multiple chains, was below 1.0 for all samples. There were no divergences in sampling. Trace plots of model sampling can be seen in figure S3 for volume and figure S6 for surface area.

### Code and computational environment

SKOOTS is operating system agnostic. All scripts were run on a computer running Ubuntu 20.04.1 LTS, an open-source Linux distribution from Canonical based on Debian. The workstation was equipped with two Nvidia A6000 graphics cards for a total of 98 GB of video memory as well as 256 GB of RAM. Many scripts were custom written in python 3.10 using open source scientific computation libraries including numpy^59^, matplotlib, scikit-learn^60^. All deep learning architectures, training logic, and much of the data transformation pipeline was written in pytorch^61^, utilizing the torchvision^61^ library. The source code may be found here: https://github.com/indzhkyulianlab/skoots.

## Acknowledgements

We would like to thank Prof. Guy Richardson for providing the block for initial FIB-SEM imaging. We also thank Dr. Stephan Kraemer (Harvard University Center for Nanoscale Systems) for their assistance in FIB-SEM imaging. This work was supported by NIH R01DC020190 (NIDCD), R01DC017166 (NIDCD), R01DC021795 (NIDCD) and R01DC017166-04S1 “Administrative Supplement to Support Collaborations to Improve the AI/ML-Readiness of NIH-Supported Data” (Office of the Director, NIH) to A.A.I. and the Speech and Speech and Hearing Bioscience and Technology Program Training grant T32 DC000038 (NIDCD). The FIB-SEM imaging was performed at the Harvard University Center for Nanoscale Systems (CNS), a member of the National Nanotechnology Coordinated Infrastructure Network (NNCI), which is supported by the National Science Foundation under NSF award 1541959. The funders had no role in study design, data collection and analysis, decision to publish, or preparation of the manuscript.

## Code availability

Code outlining SKOOTS segmentation is freely available here: https://github.com/indzhykulianlab/skoots, while training scripts, model construction, and augmentation are found here: https://github.com/indzhykulianlab/bism. SKOOTS documentation is publicly hosted here: https://skoots.readthedocs.io.

