## Supplementary Figures for "SKOOTS: Skeleton-oriented object segmentation for mitochondria"

Supplemental Figures

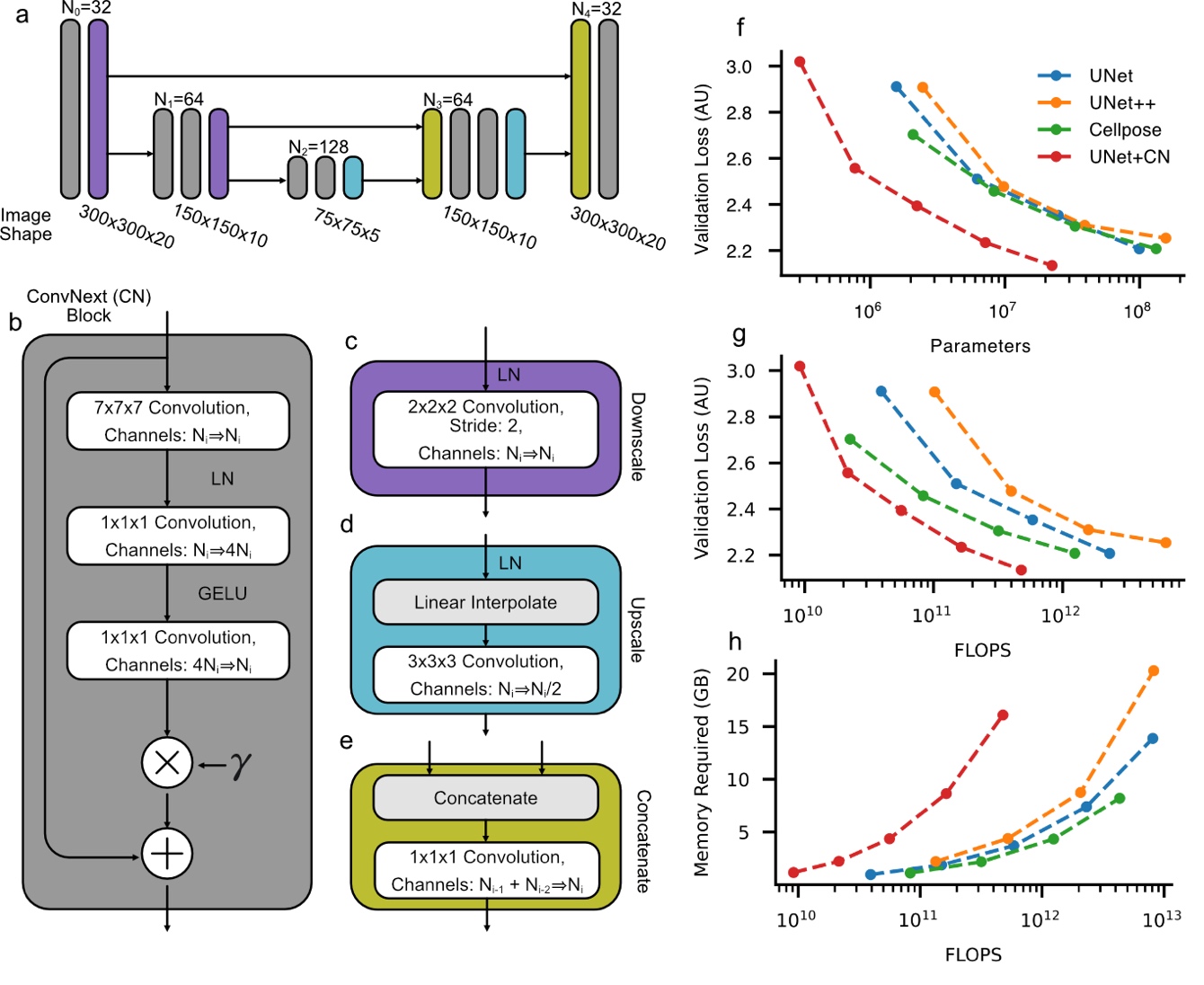

**Supplemental Figure 1. UNet Architecture with ConvNext computational blocks offers superior accuracy per floating-point operations/second (FLOPS) in a semantic segmentation task.** (**a**) Structure of a UNet-like model, composed of four major neural-network computational blocks, summarized in panels b-e, and color-coded across a-e for clarity. (**b**) a ConvNext residual block replaces the standard two stacked convolutions of UNet with a single convolution with increased kernel size (7x7x7) followed by two pointwise convolutions and a learnable scaling parameter (1/2×), (**c**) a downscale block composed by a 3D convolutional kernel with a 2x2x2 shape and a stride of 2, (**d**) an upscale block achieved by linear interpolation followed by convolution, and (**e**) a concatenation block. (**f-h**) Upon direct comparison, this architecture (**Red**) achieves superior performance semantic segmentation compared to currently reported best alternative architectures (UNet, UNet++ and Cellpose), with (**f**) a lower number of parameters, (**g**) a lower number of FLOPS, however, needs slightly more (**h**) video memory to process identically sized volumes when compared to alternatives**.**

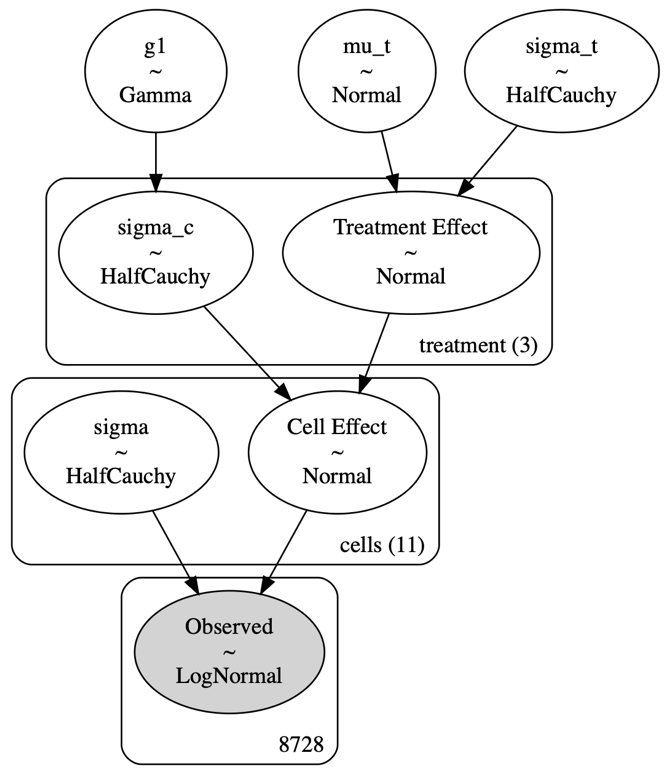

**Supplemental Figure 2**: **Bayesian hierarchical model for assessing treatment effects of aminoglycoside on mitochondria morphology.** All prior distributions were uninformative and did not significantly bias model results.

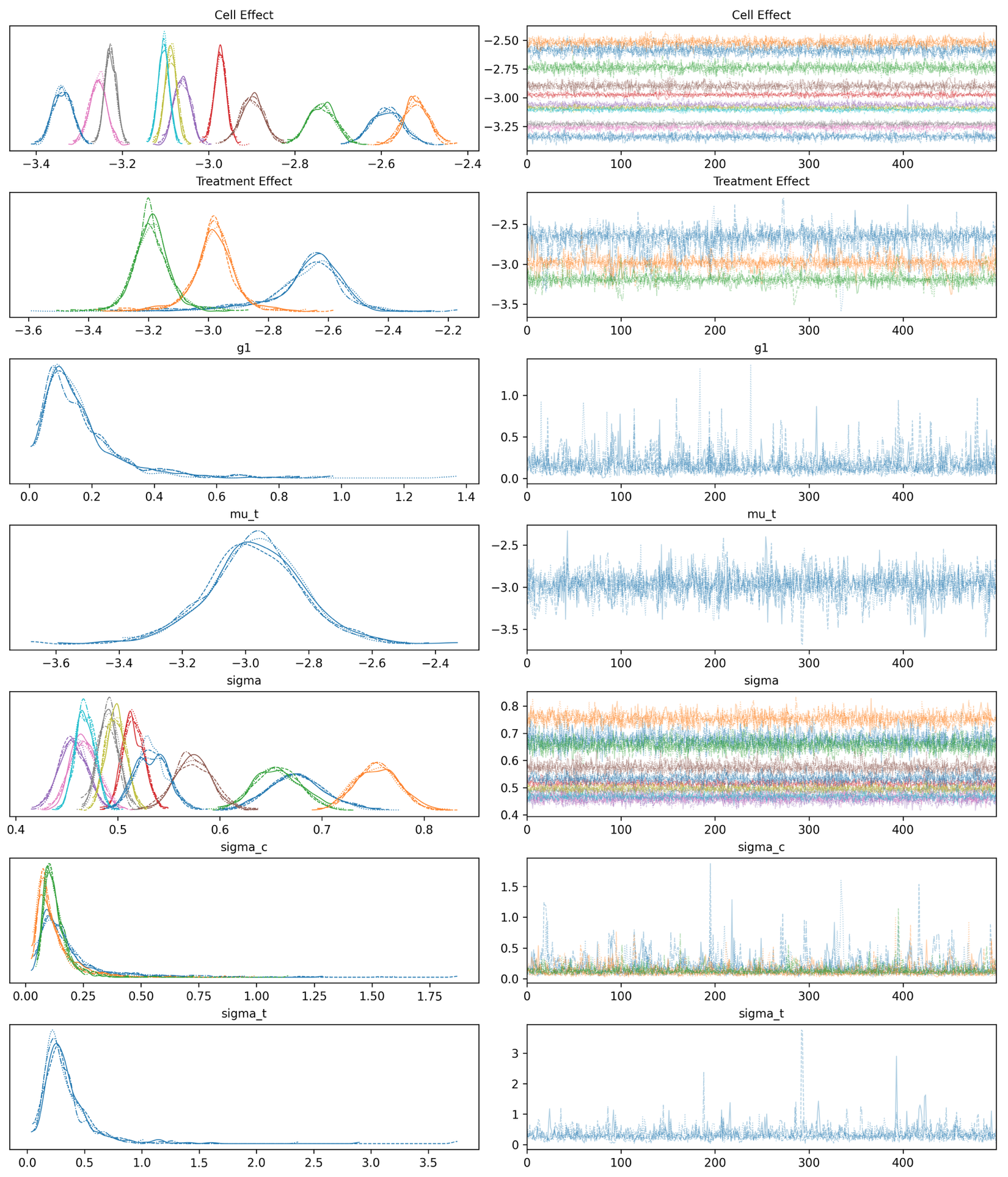

**Supplemental Figure 3: Trace plot of model sampling fit on mitochondria volume.** Four chains using the no U-turn sampler (NUTS) were sampled five hundred times to infer posterior distributions of our model. No divergences were observed and all R Hat statistics were less than one.

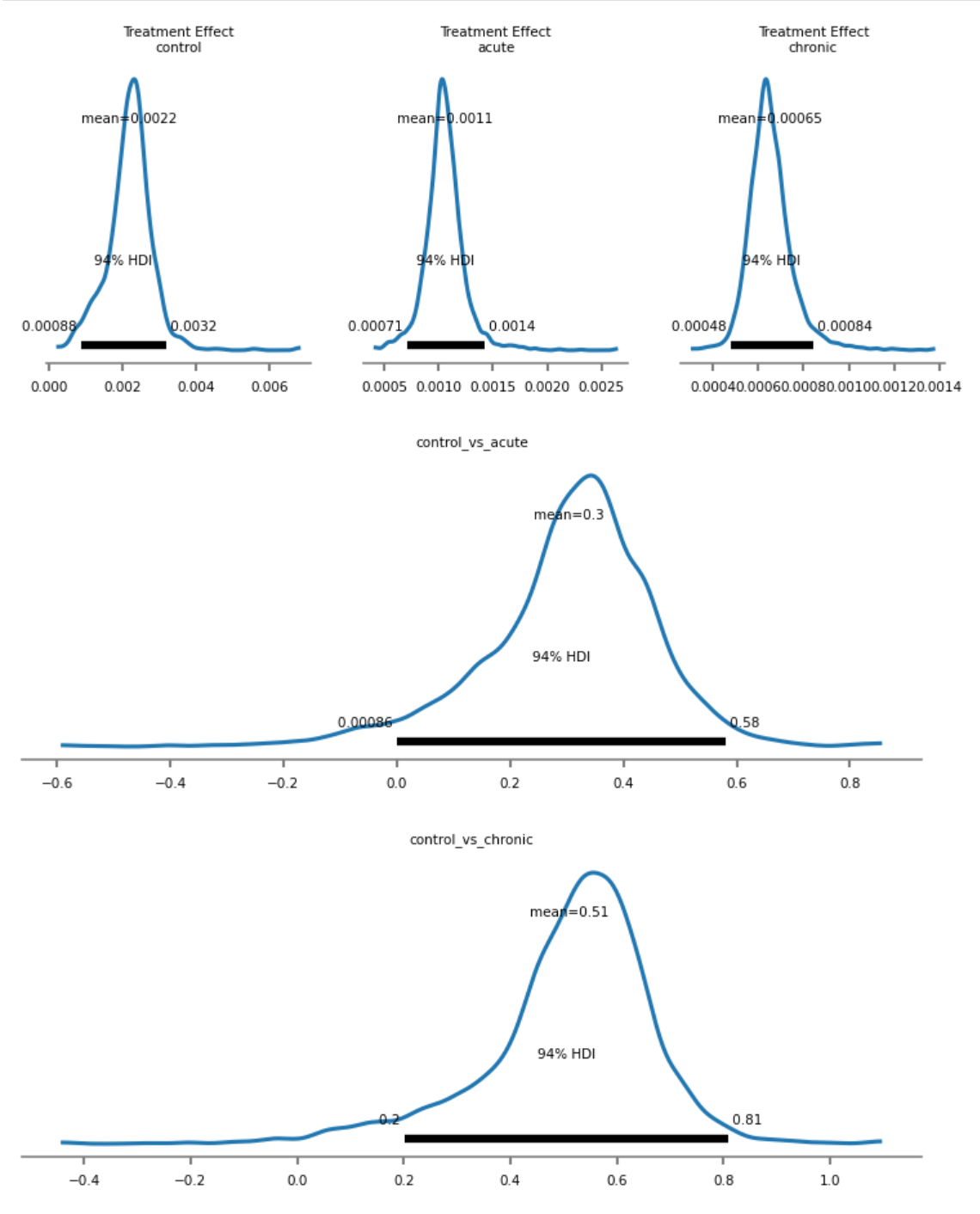

**Supplemental Figure 4:** **Posterior predictive distributions of treatment effect of aminoglycoside treatment on mitochondrial volume.** 94% credible intervals greater than zero indicate a significantly significant effect of both acute and chronically treated aminoglycoside treatment on mitochondrial volume.

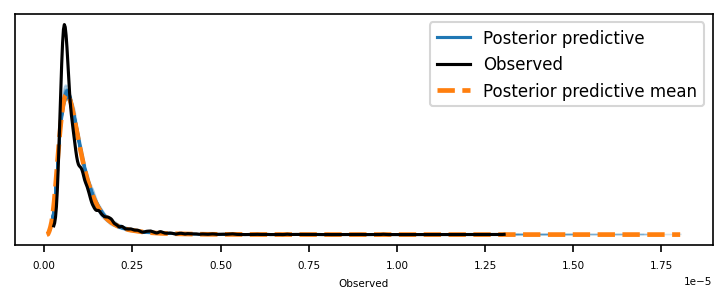

**Supplemental Figure 5** **Posterior predictive check of mitochondria volume indicates the model approximates the observed data well.**

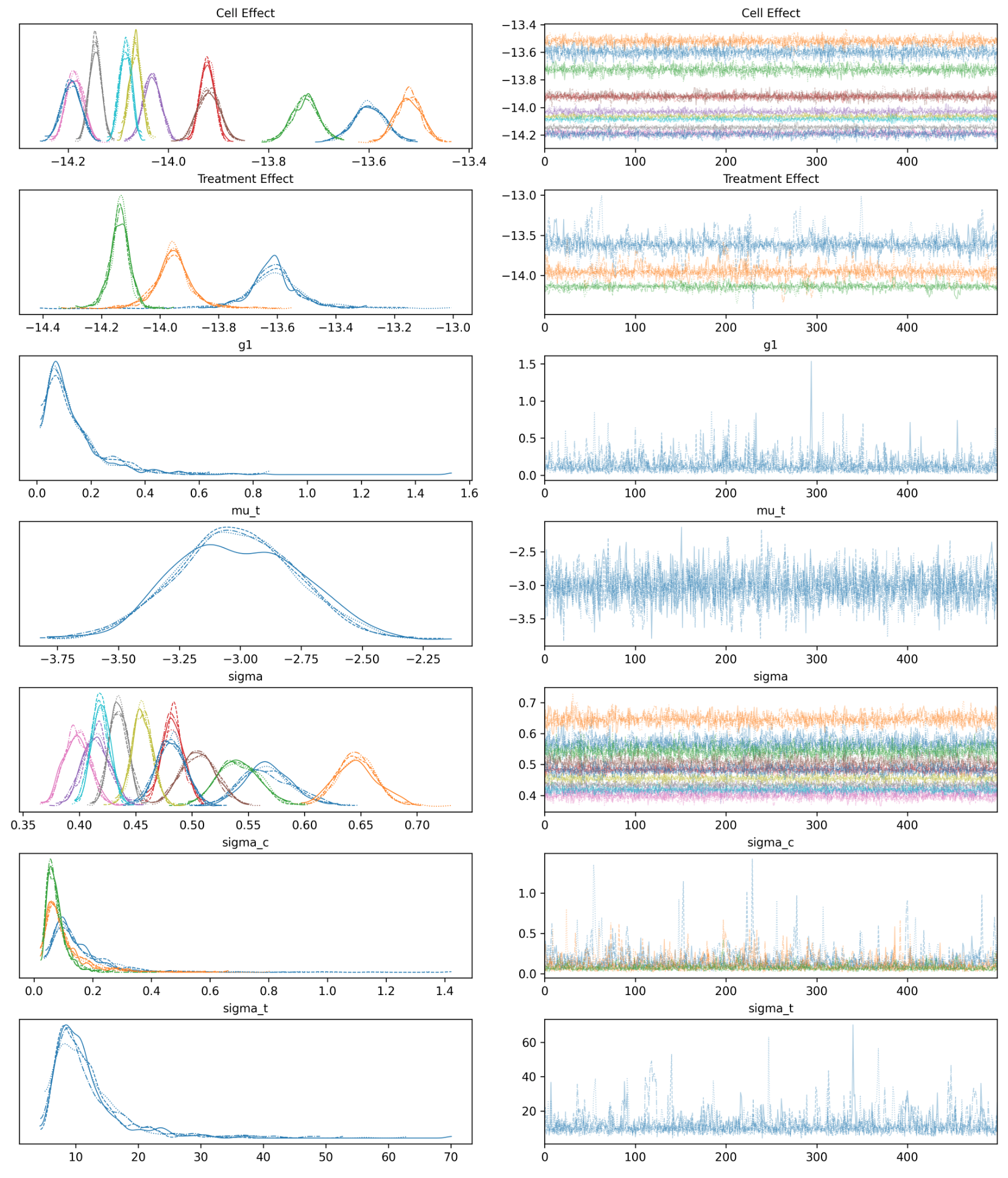

**Supplemental Figure 6: Trace plot of model sampling fit on mitochondria surface area.** Four chains using the no U-turn sampler (NUTS) were sampled five hundred times to infer posterior distributions of our model. No divergences were observed and all R Hat statistics were less than one.

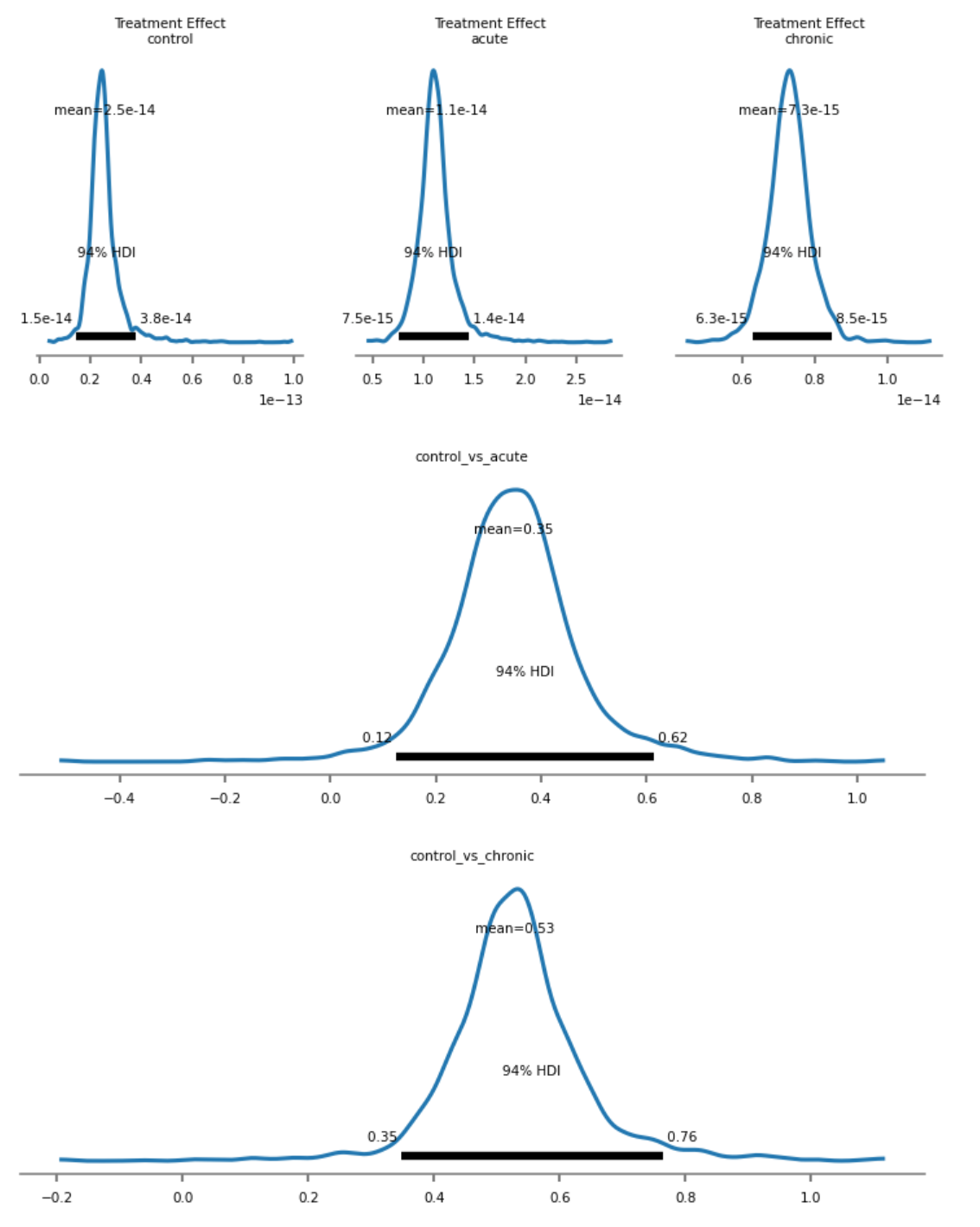

**Supplemental Figure 7 Posterior predictive distributions of treatment effect of aminoglycoside treatment on mitochondrial surface area.** 94% credible intervals greater than zero indicate a significantly significant effect of both acute and chronically treated aminoglycoside treatment on mitochondrial surface area.

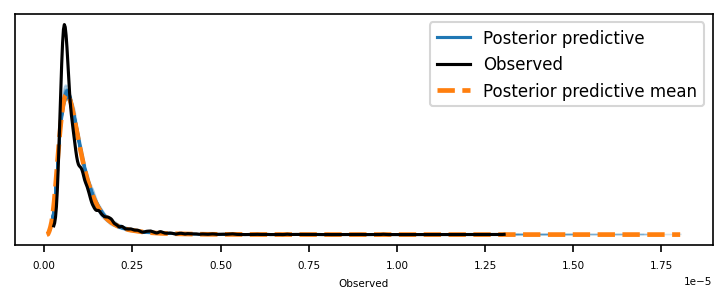

**Supplemental Figure 8 Posterior predictive check of mitochondria surface area indicates the model approximates the observed data well.**
