## Supplementary Methods for "SKOOTS: Skeleton-oriented object segmentation for mitochondria"

Supplemental Methods

INTRODUCTION

By recognizing the design patterns and protocols necessary to train a deep learning model for bio-image analysis task, we here present a new library for training segmentation models on microscopy images. This library, titled “biological image segmentation models (BISM),” builds upon the repeatability of configuration-based libraries such as pytorch connectomics, while maintaining the extensibility and verbosity of pure pytorch training. By creating a formal system for instantiating a great variety of deep learning based bioimage analysis segmentation tasks, we can construct principled comparisons of different segmentation methodologies, model architectures, or training formulae to improve the scientific rigor of these tasks. Here we present the design decisions and considerations of the library and demonstrate its usefulness by training a highly accurate model for the detection of collagen bundles in cross-section transmission electron microscopy images of the tectorial membrane.

A typical machine learning (ML) project for biological image analysis nearly always follows a supervised learning protocol. First, a parameterized model is selected and optimized to make predictions about labeled data, learning parameters that minimize prediction error. This trained model is then deployed to make the same kind of predictions for new biological images. The model architecture, image dataset, and loss functions used in training may vary for each project but are generally required to develop machine learning models. ML libraries, such as pytorch or tensorflow, are powerful and extensible but verbose, necessitating scientists to write large amounts of project-invariant code for each project. Existing high-level libraries attempt to provide a more conducive workflow for scientists by abstracting much of the programmatic scaffolding, such as pytorch lightning^1^. But by increasing abstraction, the user often relinquishes control over specific aspects of model design and training. Configuration based systems have solved this dichotomy in other domains within deep learning yet are absent for bioimage analysis. With BISM, we rectify this and present an extensible configuration based deep learning library for biomedical image analysis tasks.

DESIGN OVERVIEW

The creation of a finely tuned, accurate, and generalizable machine learning model can feel more of an art than science. Many decisions in a training pipeline can affect ultimate performance for a particular problem landscape. Much of these choices are reflected in code and are therefore difficult to track, even with proper version control. As the project evolves, training data are modified, hyperparameters may change, and deployed models may be fine-tuned from other models, making it difficult to track the recipe that led to a particular model’s good (or poor) performance. Thus, machine learning experiments are not reproducible without an understanding of the decisions made during training.

To mitigate this, we have developed BISM training pipline to be determined completely by explicitly defined configuration files which determine, with extreme granularity, the precise way to train models for various machine learning task. The configuration file is the only interface to the training pipeline, and therefore can *repeatably* map a training recipe to a model outcome with a static codebase. To achieve this, we have compartmentalized each part of a machine learning pipeline into isolated blocks which can then be selected and modified via the configuration file. The blocks of code have standardized input and outputs which allow users to freely modify core functionality simply by selecting the appropriate block in the configuration file. In this way we can dynamically create custom models, data loaders, data augmentation pipelines, training engines, and more, for dramatically different image segmentation tasks without the need for the user to modify any code. When code modifications must be made, a new block can be written and registered as a separate entity with the software. If standard inputs and outputs are honored, the registration of a new code block expands the functionality of the software while preserving backwards compatibility.

*Considerations*

With most microscopy images analyzed today being two dimensional, it is critical to support two-dimensional analysis, however modern imaging techniques have increased the accessibility of three-dimensional imaging and therefore, if possible, BISM strives to provide dimensionally agnostic implementations of all core features. To increase reproducibility of all experiments, we elect to fix the seed of all random number generators. While this has obvious implications as a specific seed may be randomly poor for one task or configuration, we believe that repeatability is paramount to good science and therefore accept this caveat. Research codebases are dynamic objects, with often poor version control practices. By decreasing the amount of code interaction, we can minimize the likelihood of code being dramatically changed and therefore enable repeatable and principled training of neural networks. Recognizing that not every model trained may have pristine handwritten logs describing an experiment, BISM strives to save as much detail as possible such that any model weights may be faithfully reproduced.

To maximize interoperability between modules of code, BISM defines a custom algorithmic type, based on a python dictionary, serving as the common language between bits of code, the aptly named *DataDict*. This *DataDict* has two key/value pairs: ‘image’ which refers to an u8int input tensor, typically of a microscopy image, and ‘masks’ referring to a u16int mask of instances where pixels/voxels of value zero are background, and positive integers representing objects in the image.

*Configuration*

Optimization of neural networks involves the determination of a multitude of hyperparameters. These hyperparameters may be values tuning the behavior of various algorithms or even the choice of algorithm itself. The agglomeration of these decisions forms the recipe of how to train a specific network and is codified into a single configuration file. We elect to use the YACS configuration system, which is based on the YAML filetype. The configuration is subdivided into the following groups: **1. System**, which describes the hardware the model will be trained on, **2. Model**, which defines the instantiation of the neural network architecture, **3. Target**, which describes the task the network will be trained on, **4. Train**, which defines all training parameters including the data, optimization algorithm, and others, and finally **5. Augmentation**, which describes the stochastic augmentation of training data during the training loop. Each configuration parameter has a default value, all of which are listed in Table 5.1.

Table SM1. BISM configuration parameters, types, and default values.

| **Group** | **Name** | **Data Type** | **Acceptable Values** | **Default Value** |
| --- | --- | --- | --- | --- |
| **SYSTEM** |  |  |  |  |
|  | NUM_GPUS | Int | > 1 | 2 |
|  | NUM_CPUS | Int | > 1 | 1 |
| **MODEL** |  |  |  |  |
|  | BACKBONE | Str | bism_unext, bism_unext2d, bism_unet, bism_unet2d, bism_unet2d_space, bism_unet3d_spade, maskrcnn_resnet50_fpn_v2 | bism_unext |
|  | MODEL | Str | spatial_embedding, lsd, generic | generic |
|  | IN_CHANNELS | Int | > 1 | 1 |
|  | OUT_CHANNELS | Int | > 1 | 3 |
|  | DIMS | List[Int] | List with length > 1 of integers > 1 | [32, 64, 128, 64, 32] |
|  | DEPTHS | List[Int] | List with length > 1 of integers > 1 | [2, 2, 2, 2, 2] |
|  | KERNEL_SIZE | Int | Either: 3, 5, 7, 9 | 7 |
|  | DROP_PATH_RATE | Float | 0.0 - 1.0 | 0.0 |
|  | LAYER_SCALE_INIT | Float | 0.0 - 1.0 | 1.0 |
|  | ACTIVATION | Str | gelu, relu, silu, sepu, tanh, sigmoid | gelu |
|  | BLOCK | Str | Block3d, block2d, unet_block3d, unet_block2d, double_spade_block3d | block3d |
|  | CONCAT_BLOCK | Str | concatconv3d, concatconv2d | concatconv3d |
|  | UPSAMPLE_BLOCK | Str | upsamplelayer3d, upsamplelayer2d | upsamplelayer3d |
|  | NORMALIZATION | Str | layernorm | layernorm |
|  | COMPILE | Bool | True, False | TRUE |
|  | OUTPUT_ACTIVATIONS | List[Str] | *List of length OUT_CHANNELS with values: gelu, relu, silu, sepu, tanh, sigmoid* | [None] |
|  | ANCHOR_SIZES | Tuple[Int] | *Tuple of length 5 with increasing integer values* | ((32,), (64,), (128,), (256,), (512,)) |
|  | ANCHOR_ASPECT_RATIOS | Tuple[Float] | *Tuple of length 3 with values 0.0-3.0* | ((0.5, 1.0, 2.0),) |
| **TRAIN** |  |  |  |  |
|  | DISTRIBUTED | Bool | True, False | TRUE |
|  | PRETRAINED_MODEL_PATH | Str | Any POSIX Path | *Nul* |
|  | TARGET | Str | lsd, affinites, mtlds, aclsd, omipose, semantic, torchvision, iadb, identity | affinites |
|  | LOSS_FN | Str | soft_dice_cldice, soft_cldice, tversky, dice, jacquard, mse, omnipose, torchvision, iade | mse |
|  | LOSS_KEYWORDS | List[Str] | *List of any keyword arguments passed to the selected loss function.* | *empty list* |
|  | LOSS_VALUES | List[Any] | *Values of any keyword arguments passed to the selected loss function.* | *empty list* |
|  | TRAIN_DATA_DIR | List[Str] | *List of any number of POSIX paths* | *empty list* |
|  | TRAIN_SAMPLE_PER_IMAGE | List[Int] | *List the same length as TRAIN_DATA_DIR with integers > 1* | [1, ] |
|  | TRAIN_BATCH_SIZE | Int | > 1 | 1 |
|  | VALIDATION_DATA_DIR | List[Str] | *List of any number of POSIX paths* | *empty list* |
|  | VALIDATION_BATCH_SIZE | Int | *List the same length as VALIDATION_DATA_DIR with integers > 1* | [1, ] |
|  | STORE_DATA_ON_GPU | Bool | True, False | FALSE |
|  | NUM_EPOCHS | Int | > 1 | 10000 |
|  | LEARING_RATE | Float | > 0.0 | 5E-04 |
|  | WEIGHT_DECAY | Float | > 0.0 | 1E-06 |
|  | OPTIMIZER | Str | adamw, adam, sgd, lion, adamax | adamw |
|  | OPTIMIZER_EPS | Float | > 0.0 | 1E-08 |
|  | SCHEDULER | Str | cosine_annealing_warm_restarts | cosine_annealing_warm_restarts |
|  | SCHEDULER_T0 | Float | > 0.0 | 10001 |
|  | MIXED_PRECISION | Bool | True, False | FALSE |
|  | N_WARMUP | Int | >= 0 | 1500 |
|  | SAVE_PATH | Str | Any POSIX Path | ./models |
|  | SAVE_INTERVAL | Int | > 0 | 100 |
|  | VALIDATION_EPOCH_SKIP | Int | > 0 | 10 |
|  | CUDNN_BENCHMARK | Bool | True, False | TRUE |
|  | AUTOGRAD_PROFILE | Bool | True, False | FALSE |
|  | AUTOGRAD_EMIT_NVTX | Bool | True, False | FALSE |
|  | AUTOGRAD_DETECT_ANOMALY | Bool | True, False | FALSE |
|  | DATALOADER_OUTPUT_DEVICE | Str | default, cpu | default |
|  | TRANSFORM_DEVICE | Str | default, cpu | default |
|  | DATASET_OUTPUT_DEVICE | Str | default, cpu | default |
|  | DATALOADER_NUM_WORKERS | Int | >= 0 | 0 |
|  | DATALOADER_PREFECTCH_FACTOR | Int | >= 0 | 0 |
| **AUGMENTATION** |  |  |  |  |
|  | CROP_WIDTH | Int | >= 1 | 300 |
|  | CROP_HEIGHT | Int | >= 1 | 300 |
|  | CROP_DEPTH | Int | >= 1 | 20 |
|  | FLIP_RATE | Float | 0.0 - 1.0 | 0.5 |
|  | BRIGHTNESS_RATE | Float | 0.0 - 1.0 | 0.4 |
|  | BRIGHTNESS_RANGE | List[Float] | [< 0.0, > 0.0] | [-0.1, 0.1] |
|  | NOISE_GAMMA | Float | >= 0.0 | 0.1 |
|  | NOISE_RATE | Float | 0.0 - 1.0 | 0.2 |
|  | CONTRAST_RATE | Float | 0.0 - 1.0 | 0.33 |
|  | CONTRAST_RANGE | List[Float] | [> 0.0, > 0.0] | [0.75, 2.0] |
|  | AFFINE_RATE | Float | 0.0 - 1.0 | 0.66 |
|  | AFFINE_YAW | List[Float] | [-180 - 180, -180 - 180] | [-180, 180] |
|  | AFFINE_SHEAR | List[Float] | [< 0, > 0] | [-7, 7] |
|  | AFFINE_SCALE | List[Float] | [> 0.0, > 0.0] | [0.85, 1.1] |
| **TARGET** |  |  |  |  |
| **TARGET.LSD** |  |  |  |  |
|  | SIGMA | Tuple[Float] | [> 0, > 0, > 0] | (8,8,8) |
|  | VOXEL_SIZE | Tuple[Float] | [> 0, > 0, > 0] | (1,1,5) |
| **TARGET.AFFINITIES** |  |  |  |  |
|  | N_ERODE | Int | >= 1 | 1 |
|  | PAD | Str | replicate | replicate |
|  | NHOOD | Int | > 1 | 1 |
| **TARGET.OMNIPOSE** |  |  |  |  |
|  | EPS | Float | > 0.0 | 1E-05 |
|  | MIN_EIKONAL_STEPS | Int | > 1 | 200 |
|  | MAX_DISTANCE | Int | > 1 | 60 |
| **TARGET.SEMANTIC** |  |  |  |  |
|  | THR | Float | 0.0 - 1.0 | 0.5 |
| **TARGET.IADB** |  |  |  |  |
|  | MASK_CHANNELS | Int | > 1 | 1 |

*Data Loading*

Images for biomedical image analysis tasks can be very large, therefore we define the atomic unit of data are an image and associated instance segmentation mask of arbitrary size. Similar images may be grouped into folders which are then loaded into memory by the software. Each image is expected to be 8bits in the TIF format and may be a 2D greyscale image, 2D color image, or 3D greyscale image. Instance segmentation masks must be a 16-bit image of the same size as the associated input image with each instance denoted by a nonzero integer, with zero reserved for background. Instance segmentation masks are linked to their associated input image by filename and the extension “labels.tif”.

An epoch is typically defined as an iteration over all pieces of data used to train a model. For computer vision, this typically means exposing each image to a model once. However, in biological image analysis, a single image may be individually too large to expose to a model in its entirety. To avoid this, BISM allows images to be grouped into separate groups which are each sampled from different number of times per epoch. Therefore, a dataset of one image sampled ten times per epoch may be represented equivalently in training to a dataset of ten smaller images sampled once per epoch. An example file structure of a training dataset is represented in Figure 5.1.

Often the most significant bottleneck in a deep learning training is the process of data loading, and their transfer to the GPU. We elect to load each image into computer memory prior to training, and when the data are sufficiently small, they may be optionally pre-loaded onto the GPU’s video memory, avoiding the bottleneck altogether.

```
training_data
├── dataset0
│   ├── very_large_training_image.tif
│   └── very_large_training_image.labels.tif
└── dataset1
    ├── another_training_image.tif
    └── another_training_image.labels.tif
```

**Figure SM1: A file structure of training data for training an instance segmentation model with BISM.** Dataset zero may be sampled more times that dataset one to ensure proper data balance in accordance to user defined configuration.

*Data Augmentation*

BISM implements multiple common geometric, and non-geometric augmentations that are dimension agnostic. Each augmentation is stochastically applied with a rate defined by the model’s configuration file. Each augmentation is dimension agnostic and may be applied on the CPU or GPU. As images in datasets may be large, and of different sizes, a mandatory crop of the image, typically at a much smaller size and at a random location, is applied. This ensures each example of a dataset is of an identical size easily digestible by the neural network. Following augmentation, each image is normalized to the dataset mean and standard deviation as normalization tends to improve the ultimate model performance^2^. The following augmentations may be applied in sequential order: Random Crop > Affine Transformation > Flip X > Flip Y > Flip Z > Adjust Brightness > Adjust Contrast > Add Noise > Normalize Image.

*Model Construction*

Convolutional neural networks with an encoder-decoder architecture are incredibly common for segmentation tasks. Popularized by Ronneberger, Fischer, Brox in 2015, the U-Net has become the de facto standard architecture for biomedical segmentation tasks ^3^. Most implementations of a U-Net have stacked double convolutions, and three to four down sampling or up sampling stages with skip connections between each down and up sampling stage. There are many variants of the U-Net architecture, some claiming to be superior to others at certain tasks. While BISM could have faithfully copied the original code for each model architecture which often explicitly define structure and (which may be advantageous for direct scientific comparison), such a practice does not allow for the flexibility to dynamically explore model architecture. Adopting a fully configuration-based approach allows the flexibility to explore the effect of novel architectural changes in a programmatic way. One such example might be the desire to explore effect of stacked *triple* convolutions in a conventional U-Net. BISM defines model architectures as only the control flow of data, excluding the operations themselves. Computations are then bundled into operational blocks with standardized signatures and selected by the user’s configuration file. BISM defines five types of operations which are bundled in the following way: upsample blocks and downsample blocks which change the resolution of the input tensor, computational blocks which perform arbitrary operations on the data while preserving spatial dimensions, regularization blocks, and finally skip connection blocks which concatenate multiple tensors into one. While BISM contains default implementations for each block, there is no enforced methodology by which each block performs its function. In an answer to the scenario before, a researcher can easily define a stacked triple convolution block and define this in the configuration file to dynamically specify the model construction and test it’s effect without completely recreating the model from scratch.

Each BISM model is defined in this way. Currently the control flow for four common architectures are implemented: U-Net, U-Net++, a Recurrent U-Net, the variant of U-Net defined by Stringer and others in Cellpose and Omipose, and a variant of U-Net employing strategies of the ConvNext architecture. An additional advantage to this system is that to switch between 2D and 3D models, only the operational blocks must be modified. When possible, each operational block will have 2D and 3D implementations. An example control flow for a generic U-Net can be seen in Figure 5.2.

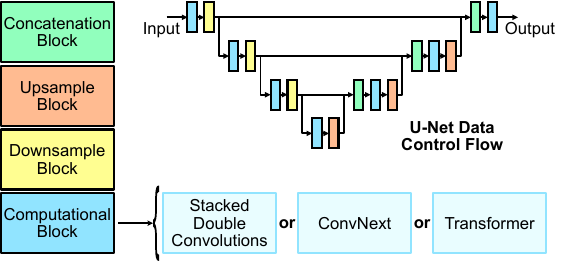

**Figure SM2. Data control flow of U-Net architecture.** BISM constructs neural network architectures by defining the flow of data through general computational operations. These operational blocks have specific function signatures, with standardized input and outputs, but BISM does not require any specific implementation. The user may therefore, via configuration, specify the computation at each part of the architecture – e.g. the user can choose either stacked double convolutions, ConvNext, or vision transformer without recreating the model in code.

In some cases, additional computation is needed to turn the output of a model into a predicted target. In many cases this is just the application of selected activation functions (relu, sigmoid, tanh, etc…), in other cases the user needs more control. Therefore, BISM optionally wraps each model with code that transforms the output of any architecture into a desired output. By default, this wrapper does nothing, however its presence allows for the flexibility of bespoke computation after the bulk of neural network computation.

*Defining a Target*

There are multiple ways to perform an instance segmentation task. Typically, a model does not directly produce an instance mask. Instead, an intermediary output is predicted by which a post-processing algorithm then creates the instance mask. For instance, a typical segmentation approach for 3D neuron reconstructions first predicts a graph of affinities, the connections between pixels of the same object, before using a watershed segmentation algorithm to generate instance masks. Another common approach, Cellpose, first predicts optical flows to an object centroid before using pixel clustering to generate resulting instance masks. In both cases, the network performs an intermediary task, the result of which is used to create the instance mask. The intermediary task is defined as the *target* of the neural network. In other scripts, these targets are calculated from the instance masks themselves ahead of time. This can reduce computation but makes geometric image augmentation difficult, as augmentation functions must therefore account for each target.

Instead, BISM defines target functions, which transform a ground truth instance mask into the neural network target after augmentation occurs. Each target function has a fixed function signature: each target function must input only a *DataDict* and a configuration file and must return the original input image tensor and the calculated target tensor. By standardizing the function signature, arbitrary target functions may be easily selected by the configuration file without the need for unique training scripts for each task.

*Training Engine*

BISM defines a training engine which executes all core functionality necessary to train a deep learning model. This includes loading data, augmenting data, instantiating optimizers, learning rate schedulers, implementing stochastic weight averaging, implementing the training loop, logging, saving, and validation. The default training engine (**Figure 5.3**) is sufficient to train neural networks for most common segmentation techniques, including affinities^4^, Cellpose^5^, Omnipose^6^, and others. Custom engines may be implemented for more complicated training pipelines, such as those required to train auto context local shape descriptors^7^.

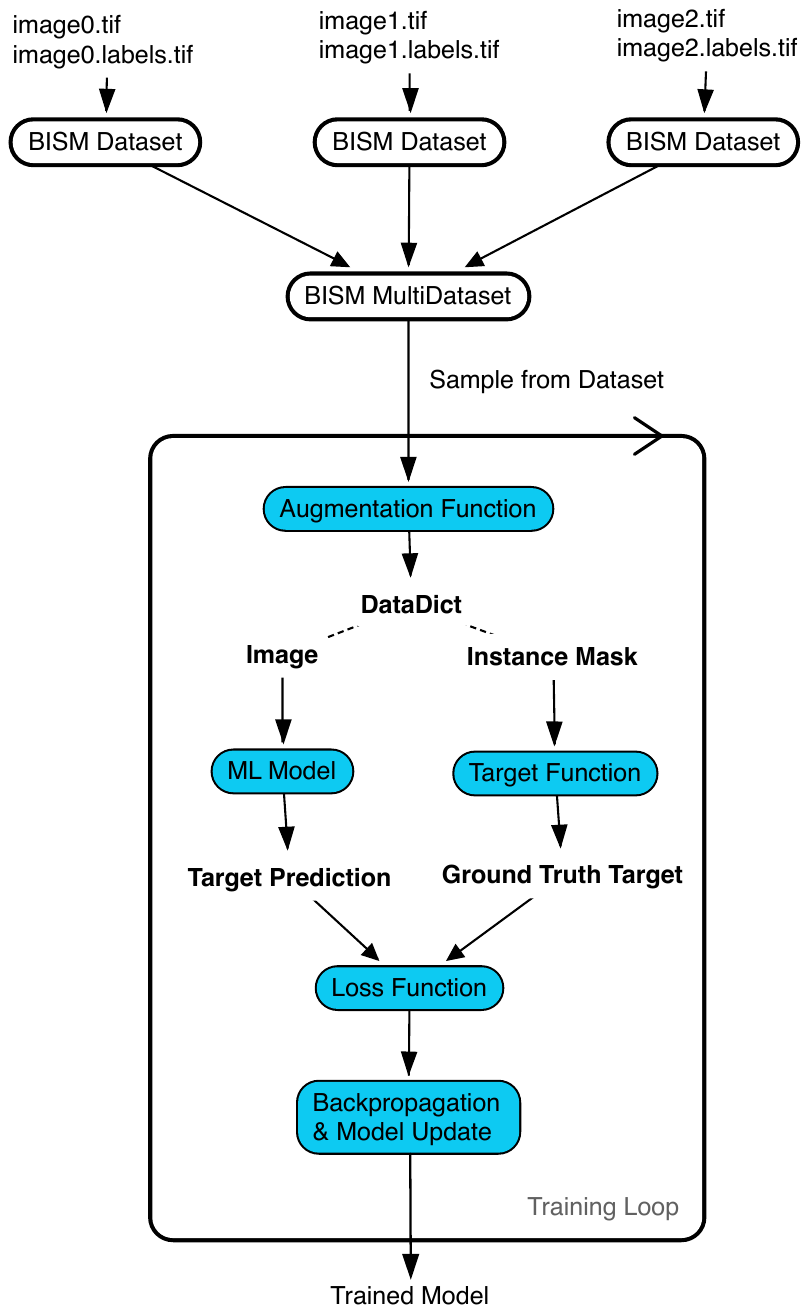

**Figure SM3 BISM Standard Training Engine.** The standard BISM training engine loads one, or many, datasets containing any number of images, or label pairs. Data are then sampled, augmented, and presented to the model until model convergence. Each step (in blue) is selected and customized via user configuration.

*Logging and Checkpointing*

BISM logs loss values and intermediate outputs of the model via the Tensorboard utility. Average training and validation loss values are recorded, along with examples of an input and output image for each training epoch. Once training has concluded, the weights and biases of the deep learning model, and optimizer, are saved. To ensure model transparency, a copy of the configuration dictating the training are saved, along with a copy of the current state of the source code, easing the ability of researchers to reproduce a trained model. Finally, while Tensorboard saves data and allows for easy model inspection, the loss values are stored in a proprietary format, difficult to access. To rectify this, each loss value is saved with the model weights. Each of the previously described configuration parameters are saved in the same file, ensuring that a researcher can always recreate a trained model, even if poor documentation or logging occurs. Finally, BISM offers a graphical user interface to inspect model checkpoint files, easing the ability to recall details on previous experiments.

*Validation*

BISM was designed to allow for rapid and principled comparison between segmentation methods for biomedical image analysis. To facilitate this, BISM includes various standard validation checks to probe model performance. Given a trained model and a reference image with ground truth annotation, BISM will calculate the average precision, true positive rate, false positive rate, false negative rate, accuracy, precision, recall, over segmentation rate and under segmentation rate.

*Extending BISM*

Due to the modular and configurable design, BISM can easily be extended to accommodate new segmentation approaches. Researchers may easily add a new variant of any configurable module and register it with the configuration system if each variant conforms to the proper function signature of that module. In some cases, training for a specific method may require bespoke functionality not offered by BISM alone. In these cases, the user may opt to add a custom training engine or choose components of the library which may be used in isolation to ease the creation of

new training paradigms.

EXPERIMENTS

*Mitochondria Segmentation by affinity prediction*

Outer hair cells are acoustic amplifiers and have considerable energetic requirements and therefore mitochondria occupy a considerable percentage of cytosolic volume. Mitochondria have little morphological homogony, and are therefore difficult to segment, however when viewed with high-resolution electron microscopy techniques, local object information, namely the object edges inferred via affinities^4^, may be predicted with sufficient accuracy to reconstruct instance masks. While suffering from known limitations^7^, affinity prediction is an efficient method for 3D object segmentation in electron microscopy. First, a deep learning model predicts a graph of the voxels representing the object, forming an sub-pixel boundary by which the watershed segmentation algorithm^8^ may use to reconstruct instance masks. Affinity prediction is an important standard technique for 3D object segmentation and therefore is included in BISM as a benchmark and reference in the development of new instance segmentation algorithms.

Instances of mitochondria in a focused ion beam scanning electron micrograph of a single murine outer hair cells was annotated to serve as training data for training an affinity detection model. Crops of the annotated dataset were randomly cropped and augmented, with affinities calculated for each instance post augmentation. Similar to above, a 3D U-Net architecture with three encoding and decoding steps with two ConvNext computational blocks^9^ at each was trained to predict X, Y, and Z affinities until convergence. To regenerate instance masks, a watershed transform was applied to the predicted affinities. For speed, multiple seed locations were detected via local maxima for each instance and watershed transform was performed in parallel. To merge over segmented masks, a subsequent agglomeration step was performed. The trained model was able to achieve accurate affinity prediction resulting in segmentation with an over segmentation rate of 1.17% on a manually annotated validation set.

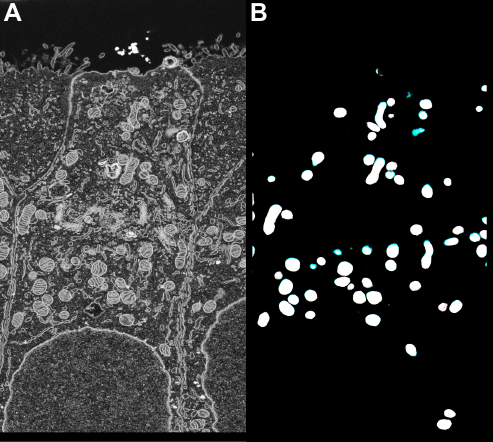

**Figure SM4: BISM affinity prediction.** A) A neural network was input a murine outer hair cell imaged with focused ion beam scanning electron microscopy and produces predicted affinity graphs (B) of mitochondria. Positive affinities in x, y, and z are represented as red, green, and blue voxel values respectively. White voxels are connected in all cardinal directions.

1.	Sawarkar K. *Deep Learning with PyTorch Lightning: Swiftly Build High-performance Artificial Intelligence (AI) Models Using Python.* Packt Publishing Ltd; 2022.

2.	Zheng Q, Yang M, Yang J, Zhang Q, Zhang X. Improvement of generalization ability of deep CNN via implicit regularization in two-stage training process. *IEEE Access.* 2018;6:15844-15869.

3.	Ronneberger O, Fischer P, Brox T. U-net: Convolutional networks for biomedical image segmentation. Paper presented at: International Conference on Medical image computing and computer-assisted intervention2015.

4.	Funke J, Tschopp F, Grisaitis W, et al. Large scale image segmentation with structured loss based deep learning for connectome reconstruction. *IEEE transactions on pattern analysis and machine intelligence.* 2018;41(7):1669-1680.

5.	Stringer C, Wang T, Michaelos M, Pachitariu M. Cellpose: a generalist algorithm for cellular segmentation. *Nature methods.* 2021;18(1):100-106.

6.	Cutler KJ, Stringer C, Lo TW, et al. Omnipose: a high-precision morphology-independent solution for bacterial cell segmentation. *Nature methods.* 2022:1-11.

7.	Sheridan A, Nguyen T, Deb D, et al. Local Shape Descriptors for Neuron Segmentation. *bioRxiv.* 2021:2021.2001.2018.427039.

8.	Angulo J, Jeulin D. Stochastic watershed segmentation. Paper presented at: ISMM (1)2007.

9.	Liu Z, Mao H, Wu C-Y, Feichtenhofer C, Darrell T, Xie S. A convnet for the 2020s. Paper presented at: Proceedings of the IEEE/CVF Conference on Computer Vision and Pattern Recognition2022.

10.	Kaplan J, McCandlish S, Henighan T, et al. Scaling laws for neural language models. *arXiv preprint arXiv:200108361.* 2020.

11.	Zhang Z. Improved adam optimizer for deep neural networks. Paper presented at: 2018 IEEE/ACM 26th International Symposium on Quality of Service (IWQoS)2018.

12.	Lin Z, Wei D, Lichtman J, Pfister H. PyTorch connectomics: a scalable and flexible segmentation framework for EM connectomics. *arXiv preprint arXiv:211205754.* 2021.

13.	Yadan O. Hydra-a framework for elegantly configuring complex applications. In: Github San Francisco, CA, USA; 2019.
